# Myeloid States Predict Fate of Premalignant Lesions

**DOI:** 10.1101/2025.11.02.686127

**Authors:** Thomas D. Madsen, Dongya Jia, Desu Chen, Maria O. Hernandez, Sarah M. Hammoudeh, Edmund Cauley, Marco Heydecker, Desiree Tillo, Madeline Wong, Emily Chen, Salma Abu-Elnaj, Ross Lake, Noemi Kedei, Gregoire Altan-Bonnet, Weiye Wang, Roberto Weigert

**Affiliations:** Laboratory of Cellular and Molecular Biology, Center for Cancer Research, National Cancer Institute, National Institutes of Health, Bethesda, MD, USA; Copenhagen Center for Glycomics, University of Copenhagen, Department for Cellular and Molecular Medicine, Copenhagen, Denmark; Immunodynamics Group, Laboratory of Integrative Cancer Immunology, Center for Cancer Research, National Cancer Institute, Bethesda, MD, USA; Spatial Imaging Technology Resource, Center for Cancer Research, National Cancer Institute, National Institutes of Health, Bethesda, MD, USA; Advanced Biomedical Computational Sciences, Frederick National Laboratory for Cancer Research, Frederick, MD, 21702, USA; CCR Genomics Core, Center for Cancer Research, National Cancer Institute, National Institutes of Health, Bethesda, MD 20892, USA; Laboratory of Cancer Biology and Genetics, Center for Cancer Research, National Cancer Institute, National Institutes of Health, Bethesda, MD, USA

**Keywords:** Premalignant lesions, Spontaneous regression, Head and neck cancer, Intravital microscopy, Spatial Transcriptomics, Myeloid Niche, Cxcl9, Cxcl10, Tumor metabolism, M2 macrophage, Tumor-Associated Macrophage, Immune checkpoint inhibitor, Tumor microenvironment

## Abstract

The spontaneous regression of cancer lesions illustrates the power of immune surveillance; yet these transient events have largely escaped systematic analysis. Using longitudinal intravital microscopy in a carcinogen-induced model of head and neck cancer, we followed premalignant lesions within the same animals for 24 weeks at single-cell resolution. This strategy uncovered three trajectories: progression, stability, or regression, and enabled direct analysis of immune dynamics underlying each fate. Lesion outcome was determined by the spatial organization of myeloid-derived antigen-presenting cells: regressing lesions were characterized by dense clusters of myeloid-derived cells associated with CXCL9^+^/CXCL10^+^ expression and T cell recruitment, whereas progressing lesions displayed a scattered infiltration of these cells. Remarkably, transient myeloid clusters arose prior to any detectable lesion formation and marked regions that would later develop into premalignant lesions. These findings identify spatiotemporal myeloid organization as an early determinant of tumor fate and provide a mechanistic framework for predicting and intercepting cancer at its inception.

**Summary Sentence:** Early myeloid architecture dictates cancer fate: dense CXCL9⁺/CXCL10⁺ clusters with T-cell enrichment accompany regression, whereas sparse infiltration predicts progression. Transient pre-lesional myeloid clusters emerge at future tumor sites, revealing immune organization as an early determinant of malignancy.

## Introduction

Cancer development reflects a dynamic interplay between environmental exposures and somatic mutations, and remains a major global health burden, with incidence and mortality continuing to rise worldwide^1^. Although malignant progression is often viewed as unidirectional, spontaneous regression of cancer lesions has been documented clinically, including in head and neck squamous cell carcinoma (HNSCC)^2–4^. These rare events demonstrate that the body possesses an intrinsic capacity to eliminate cancer without therapeutic intervention. However, because spontaneous regression is transient and unpredictable, it is rarely accessible for systematic study, limiting mechanistic insight into how immune responses succeed or fail during early tumorigenesis. Immune checkpoint blockade (ICB) illustrates the therapeutic potential of restoring immune control by blocking inhibitory pathways such as PD1/PD-L1 and CTLA4^5^. Despite its clinical impact, only a minority of patients derive durable benefits^6,7^, and immune-related toxicity remains a major limitation^8^. While the absence of T-cell infiltration characterizes immunological “cold” tumors, T-cell-rich lesions often also fail to regress, indicating that the immune cell presence alone is insufficient to predict outcome^9,10^. Increasing evidence highlights a critical role for the innate immune compartment, showing that spatially organized multicellular niches composed of antigen-presenting cells and T cells predict favorable outcomes across several cancers^11–14^. These findings underscore that immune function is shaped not only by cell identity but also by spatial organization within the tumor microenvironment.

Despite these advances, most mechanistic insights into tumor–immune interactions derive from established tumors analyzed at single time points, frequently using xenograft or genetically engineered mouse models. Such systems capture late-stage disease, after lesion fate has already been determined, and therefore cannot resolve how immune dynamics influence whether premalignant lesions regress, stabilize, or progress. Addressing this gap requires approaches capable of tracking immune behavior longitudinally and at cellular resolution during the earliest phases of neoplastic evolution.

Longitudinal intravital microscopy provides a unique opportunity to meet this challenge by enabling repeated imaging of living tissues at single-cell resolution over time. This approach has yielded fundamental insight into invasion^15–18^, metastasis^19,20^, tumor metabolism^21^, and immune responses in established tumors^17,22–26^. However, its application to premalignant disease has been limited. To overcome this barrier, we adapted intravital microscopy to follow the evolving microenvironment of the oral epithelium in a carcinogen model of HNSCC. Administration of 4-nitroquinoline-1-oxide (4NQO) induces stepwise transformation of the tongue epithelium^27–29^ and closely mirrors the mutational landscape of human disease. Importantly, this model shows partial responsiveness to PD-1 blockade^30^ and features a slow and stochastic lesion onset, which enables longitudinal analysis of early pathological stages^31^. Using this platform, we recently observed that only a subset of 4NQO-induced lesions progress to carcinoma, whereas others persist or undergo spontaneous regression within the same animal (Wang et al., man in prep).

Here, we combine longitudinal intravital microscopy with correlative multiplex immunostaining and spatial transcriptomics to define how myeloid cells behave during early neoplastic evolution and how their spatial organization influences outcome. We show that antigen-presenting myeloid cells dominate the immune infiltrate of premalignant lesions and that their spatial organization, rather than their mere presence, distinguishes regressing from progressing trajectories. By resolving immune dynamics before, during, and after lesion emergence, this work identifies spatiotemporal myeloid architecture as a critical determinant of tumor fate and establishes a framework for predicting and intercepting cancer at its inception.

## Results

### Longitudinal intravital imaging reveals three trajectories of early lesion fate

To longitudinally track premalignant lesion development *in vivo*, we performed biweekly intravital microscopy of the ventral tongue in LysM-cre; mT/mG dual reporter mice^32^ treated with 4NQO (Fig. 1A, see method section). This approach enabled simultaneous visualization of myeloid cells (GFP⁺), epithelial architecture (TdTomato⁺), and the collagen-rich stromal matrix via second harmonic generation (SHG)^33^, allowing repeated imaging of the same tissue regions over the course of carcinogenesis.

**Figure 1.**
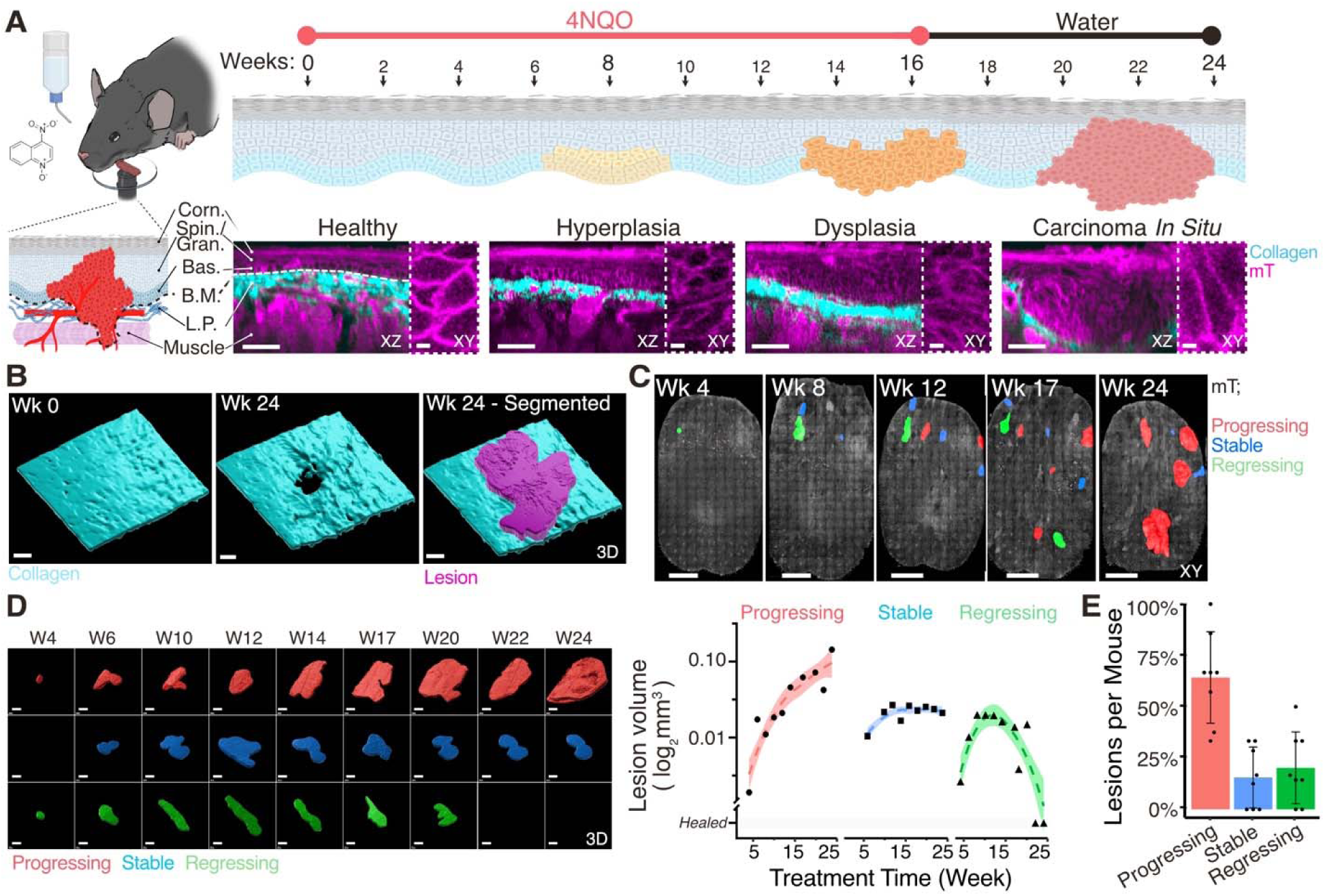
Longitudinal intravital microscopy reveals distinct trajectories of premalignant lesion fate. Eight LysM-Cre; mT/mG mice expressing membrane-bound TdTomato (mT) in all cells and GFP (mG) in myeloid cells were treated with 50 µg/mL 4NQO in drinking water for 16 weeks, followed by an 8-week recovery period. The rostral two-thirds of the ventral tongue were imaged biweekly by intravital two-photon microscopy to a depth of 150 µm. **(A)** Schematic of the longitudinal imaging workflow and tongue epithelial architecture. Representative optical sections illustrate epithelial morphology at different stages of premalignant progression mT channel gamma is set to 1.5 to outline epithelial morphology. Scale bar, 50 µm. Inserts show the morphology of the basal epithelial layer. Scale bar, 5 µm **(B).** Three-dimensional segmentation of the lamina propria (collagen, SHG)) and lesion epithelium, highlighting collagen displacement and altered cell morphology. Scale bar, 100 µm. **(C)** Whole-tongue images showing the spatial distribution of segmented lesions across the ventral surface. Colored outlines denote lesions with different growth behaviors. Scale bar, 1 mm. **(D)** Temporal 3D segmentation of representative lesions illustrating progression, regression, or stability. Loess fit of log₂-transformed lesion volumes over time shows exponential growth in progressing lesions, and minimal growth or regression in the others. Scale bar, 200 µm. **(E)** Summary of lesion outcomes across all mice. Fifty-two lesions (∼500 imaging time points) were recorded: 28 progressed, 8 stabilized, and 11 regressed (6 partial, 5 complete). On average, each mouse developed ∼4 progressing, 1-2 regressing, and 1 stable lesion. Lesions that evolved into benign papillomas were excluded from analysis. Corn, corneal layer; Spin, spinous layer; Gran, granular layer; Bas, basal layer; BM, basement membrane; LP, lamina propria.

Premalignant lesions were identified based on localized epithelial morphology alterations and associated collagen deformation (Fig. 1B–D). Three-dimensional segmentation of the epithelial compartment enabled quantitative tracking of lesion volumes over time. (Fig. S1A-D, see Method Section). Lesions were segregated into three categories: progressing (64 ± 22%, mean ± S.D.), regressing (20 ± 17% mean ± S.D.; 11% partial / 9% complete remission), and stable (16 ± 15% mean ± S.D.) (Fig. 1E). Progressing lesions exhibited an average volumetric growth rate of 24 ± 10% (mean ± S.D.) per week (Fig. S1E). Each animal developed on average 3–5 progressing lesions, 1–2 regressing lesions, and one stable lesion. The coexistence of all three trajectories within the same animal ruled out systemic factors and framed a platform to dissect the local immune determinants of lesion fate. Note that stability was defined relative to the 24-week observation window, and lesions classified as stable may have subsequently regressed or progressed beyond the duration of the study. These proportions are consistent with observations from other mouse strains (Wang et al., man. in prep.), which reported comparable distributions across different mouse strains, underscoring the robustness of this phenomenon.

Together, these data establish a longitudinal in vivo framework that captures divergent early lesion trajectories and enables interrogation of the local immune and tissue dynamics governing premalignant fate decisions

### Infiltrating myeloid behavior and tumor metabolism predict lesion fate

In the healthy tongue, myeloid cells were sparsely distributed across distinct tissue compartments, with dendritic cell (DC)–like cells present in the epithelium, macrophage-like cells in the lamina propria, and neutrophil-like cells confined to blood vessels (Fig. S2A–C). A subset of perivascular GFP⁺ cells lacking CD45, CD68, MHCII, and Langerin expression remained immobile throughout 24 weeks of imaging (Fig. S2D), consistent with partial neuronal expression previously reported in LysM-Cre mice^34^. Following 4NQO treatment, a marked increase in myeloid cells was observed throughout the tongue (Fig. 2A, S3A). Control mice imaged under identical conditions showed neither lesion formation nor myeloid recruitment, confirming that recruitment was driven by the 4NQO treatment rather than the imaging procedure.

**Figure 2.**
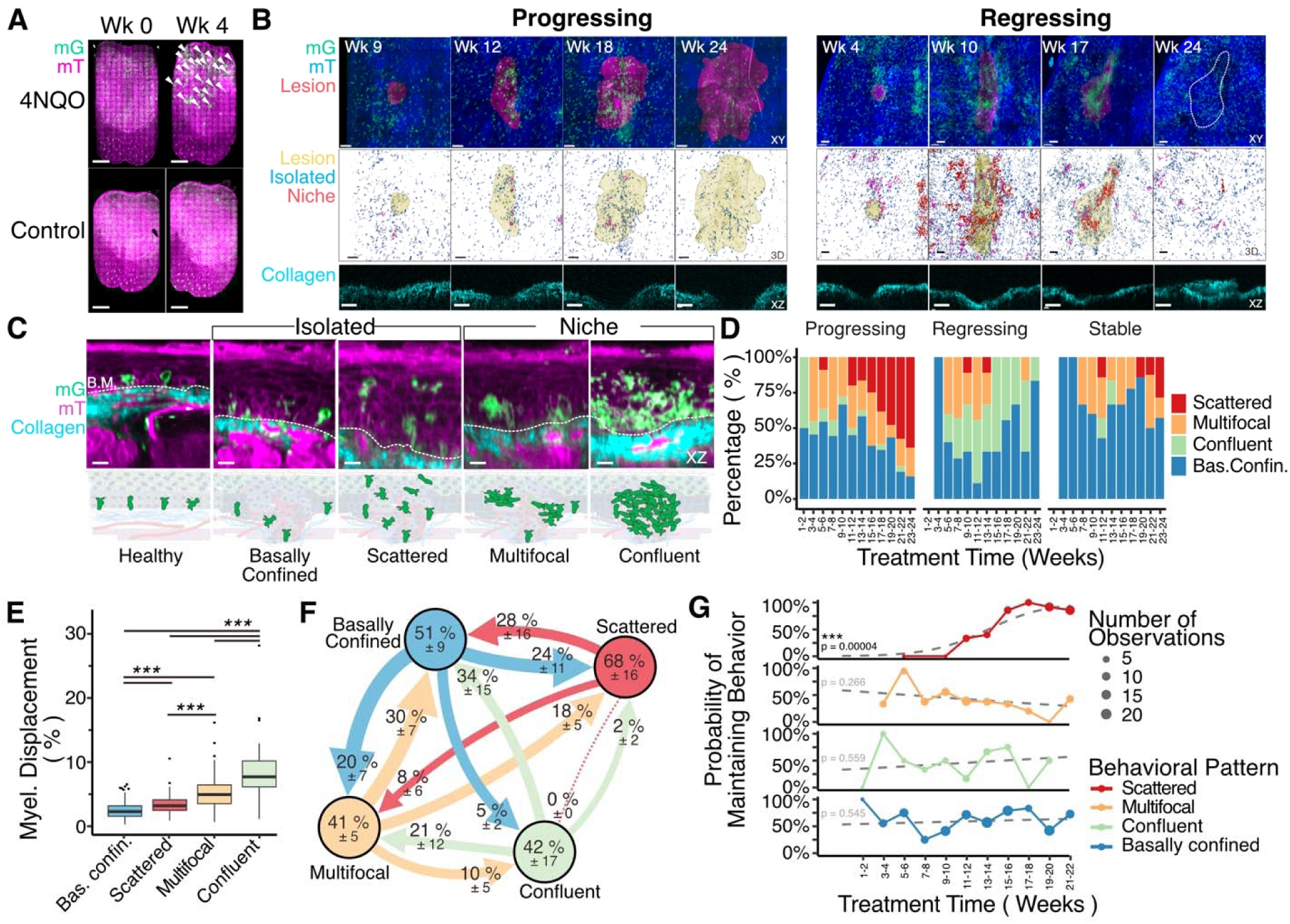
Transient myeloid behavioral states predict the progression or regression of premalignant lesions. Myeloid cells (GFP⁺) were imaged in the tongues of LysM-Cre; mT/mG mice during the 24-week 4NQO treatment regimen described in Fig. 1. (A) Whole-tongue scans before and after 4 weeks of 4NQO treatment show increased myeloid infiltration following carcinogen exposure (arrowheads). Vehicle-treated control mice showed a stable baseline of myeloid cells at both time points (see also Fig. S2). Scale bar, 1 mm. (B) Sum projections of lesion epithelium illustrating myeloid infiltration in a progressing lesion (left) and a regressing lesion (right). Middle panels show 3D segmentation of isolated (< 5000 µm³; blue) and niche-forming (> 5000 µm³; red) myeloid cells reveals larger and more frequent niches in regressing lesions. Bottom panels show cross-sections of the collagen layer, highlighting collagen restoration following lesion regression. Scale bar, 100 µm. (C) Cross-sectional views illustrating the four principal intratumoral myeloid organizational states: basally confined, scattered, multifocal, and confluents. Scale bar, 100 µm. (D) Predominant myeloid organization observed over time, stratified by lesion outcome. (E) Quantification of intralesional myeloid displacement (fraction of lesion volume occupied by myeloid cells) stratified by organizational state. Statistical comparisons were performed using the Kruskal–Wallis test with pairwise Wilcoxon post hoc testing. (F) Transition probability between myeloid organizational states across consecutive imaging sessions (2-weeks intervals). Values represent mean ± SEM (n = 6–67 transitions per mouse, 8 mice total). (G) Probability of maintaining each myeloid organization as a function of treatment time. Associations between behavioral maintenance and lesion age were tested using the Cochran–Armitage trend test. (G) Cross-sectional NADH fluorescence images from the same lesions shown in (A), exhibiting early NADH decline preceding myeloid infiltration in progressing lesions and coincident with infiltration in regressing lesions. Representative images are shown from three progressing and three regressing lesions. Scale bar, 100 µm. mT/mG: Membrane-bound TdTomato/GFP.

Within premalignant lesions, myeloid cells displayed distinct spatial organization patterns that correlated with lesion trajectory (Fig. 2B). In progressing lesions, infiltrating myeloid cells were predominantly isolated and dispersed, whereas regressing lesions were characterized by multicellular immune niches. To capture these organizational states, intralesional myeloid behavior was classified into four categories reflecting increasing cellular interaction and density: basally confined, scattered, multifocal, and confluent (Fig. 2C). Basally confined myeloid cells aligned along the basement membrane, resembling the organization of healthy tissue. Scattered cells infiltrated into the suprabasal layers as isolated cells, losing contact with the basement membrane (Supplementary Video 1. Multifocal lesions contained multiple small clusters, while confluent lesions exhibited large, continuous aggregates occupying a substantial portion of the lesion epithelium (Supplementary Video 2).

At early stages, lesions of all trajectories predominantly exhibited basally confined or transient multifocal myeloid organizations (Fig. 2D, S3B). Over time, diverging patterns emerged. Progressing lesions increasingly adopted a scattered organization, with cells displaying dendritic morphology and active antigen-scavenging behavior. In contrast, regressing lesions formed confluent myeloid niches composed of cells containing endocytosed TdTomato signal and a foamy cytoplasm, consistent with increased phagocytic activity (Fig. S3C). Stable lesions largely retained basally confined or multifocal patterns. Intralesional myeloid displacement, defined as the fraction of lesion volume occupied by myeloid cells, increased monotonically from basally confined through scattered, multifocal, to confluent patterns (Fig. 2E, S3D–E).

Longitudinal tracking revealed that myeloid organization was dynamic within individual lesions (Fig. 2F). The scattered pattern was the most stable, persisting across consecutive imaging sessions in 68 ± 16% (mean ± S.E.M.) of cases, whereas multifocal and confluent organizations were more transient, changing state in over half of sequential timepoints. Notably, multifocal patterns more frequently reverted to basally confined organization than progressed toward confluent infiltrates. Analysis of temporal stability showed that the probability of maintaining a scattered configuration between two consecutive timepoints increased with lesion age, whereas multifocal organization became progressively less stable over time (Fig. 2G).

Because tumor metabolism is tightly linked to immune behavior^22,35^, we next examined endogenous NADH fluorescence as a real-time, label-free indicator of epithelial metabolic state^36^ (Wang et al man in prep). This analysis revealed trajectory-dependent metabolic patterns (Fig. 3A-B). Many progressing lesions exhibited an early and sustained decline in NADH intensity, often preceding myeloid infiltration. In contrast, regressing lesions maintained NADH levels until the emergence of confluent myeloid niches, at which point a transient NADH decrease was occasionally observed. Stable lesions largely maintained consistent NADH levels throughout their lifetime. High-resolution imaging confirmed that loss of NADH signal occurred within intact epithelial cells, as TdTomato membrane labeling remained preserved (Fig. 3C). Stratification of myeloid behavior by metabolic state revealed a strong association between low intralesional NADH and scattered myeloid organization (Fig. 3D). Temporal analysis across individual lesions showed variable ordering of NADH decline and myeloid scattering, with no consistent sequence observed (Fig. 3E). Thus, while NADH loss and scattered myeloid behavior were tightly correlated with lesion progression, their temporal relationship varied across lesions.

**Figure 3.**
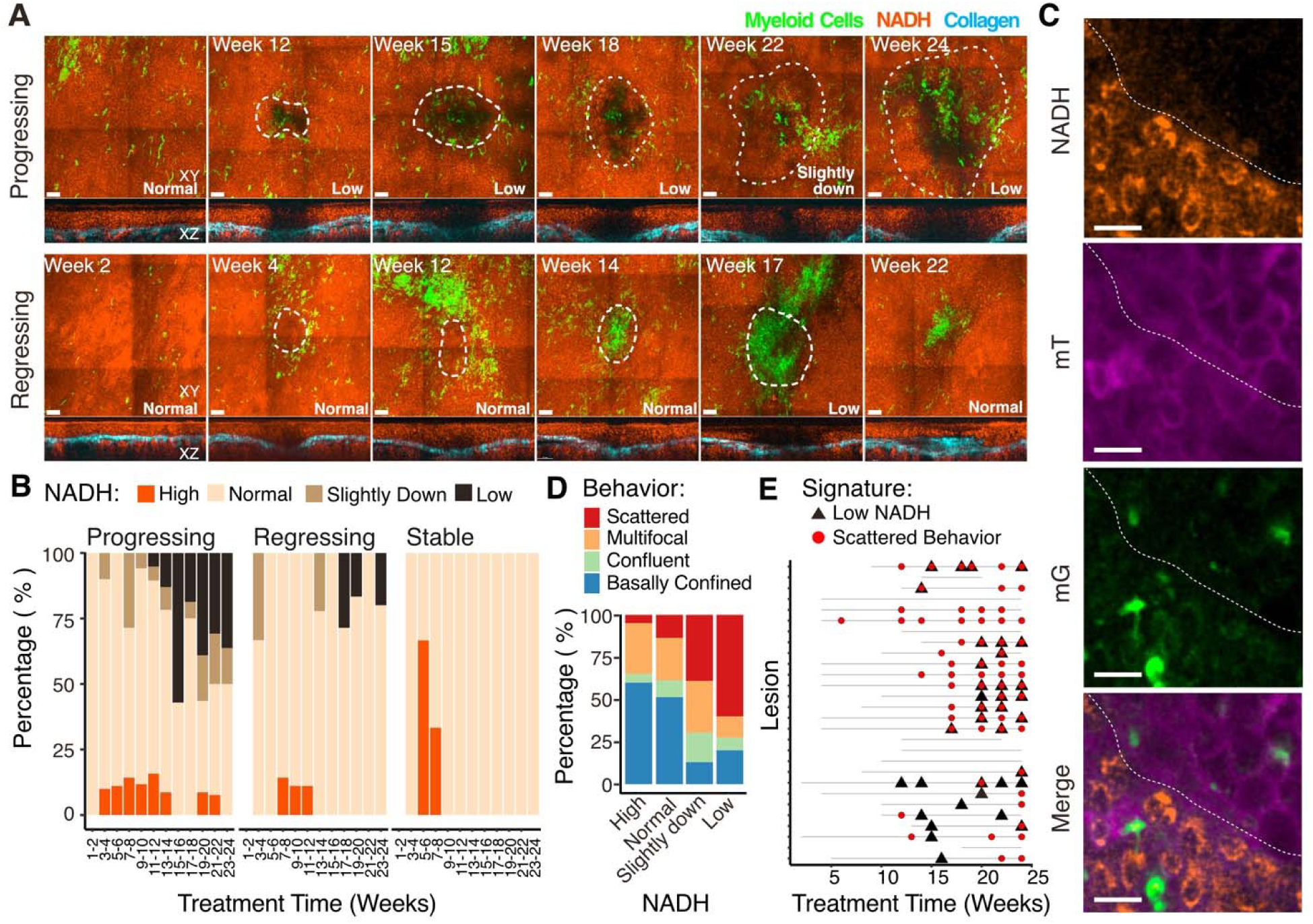
A metabolic switch accompanies scattered myeloid organizations and is predictive for lesion trajectory. Endogenous NADH fluorescence was imaged longitudinally as a label-free readout of epithelial metabolic state. **(A)** Sum-projections of the epithelial layer showing loss of NADH within lesions relative to myeloid infiltration (XY view). Dashed lines indicate lesion boundaries. Side view (XZ) shows NADH signal and collagen signals only. **(B)** Quantification of NADH level over time stratified by lesion trajectories. **(C)** High magnification optical slices at the boundary between high- and low-NADH regions within a progressing lesion. Membrane Tomato labeling remains intact in low-NADH epithelial cells. **(D)** Fraction of lesion time points exhibiting each myeloid organization stratified by the NADH state. **(E)** Temporal relationship between scattered myeloid cell organization and low-NADH state across individual lesions. mT/mG: Membrane-bound TdTomato/GFP.

Together, these data identify two early features associated with premalignant lesion progression: a transition to scattered myeloid organization and a decline in epithelial NADH levels. Both features provide real-time, label-free indicators of lesion trajectory and precede overt malignant transformation.

### Myeloid-derived dendritic cells are the primary infiltrating myeloid cell type

To define the immune cell types underlying the functional states of confluent and scattered infiltrates associated with regression and progression, respectively, we developed a correlative workflow that combined longitudinal intravital microscopy (IVM) with endpoint cyclic multiplex immunofluorescence (mIF) based on the Iterative Bleaching Extends Multiplexity (IBEX) protocol^37^. Tongues were embedded in an orientation to section the ventral epithelium as intact sheets that enables spatial registration between intravital imaging volumes and histological sections and allows direct cell-to-cell correlation between intravital and histological datasets (Fig. 4A). This approach was applied to mice at weeks 8, 16, and 24, as well as untreated controls, capturing lesions across multiple stages of premalignant evolution.

**Figure 4.**
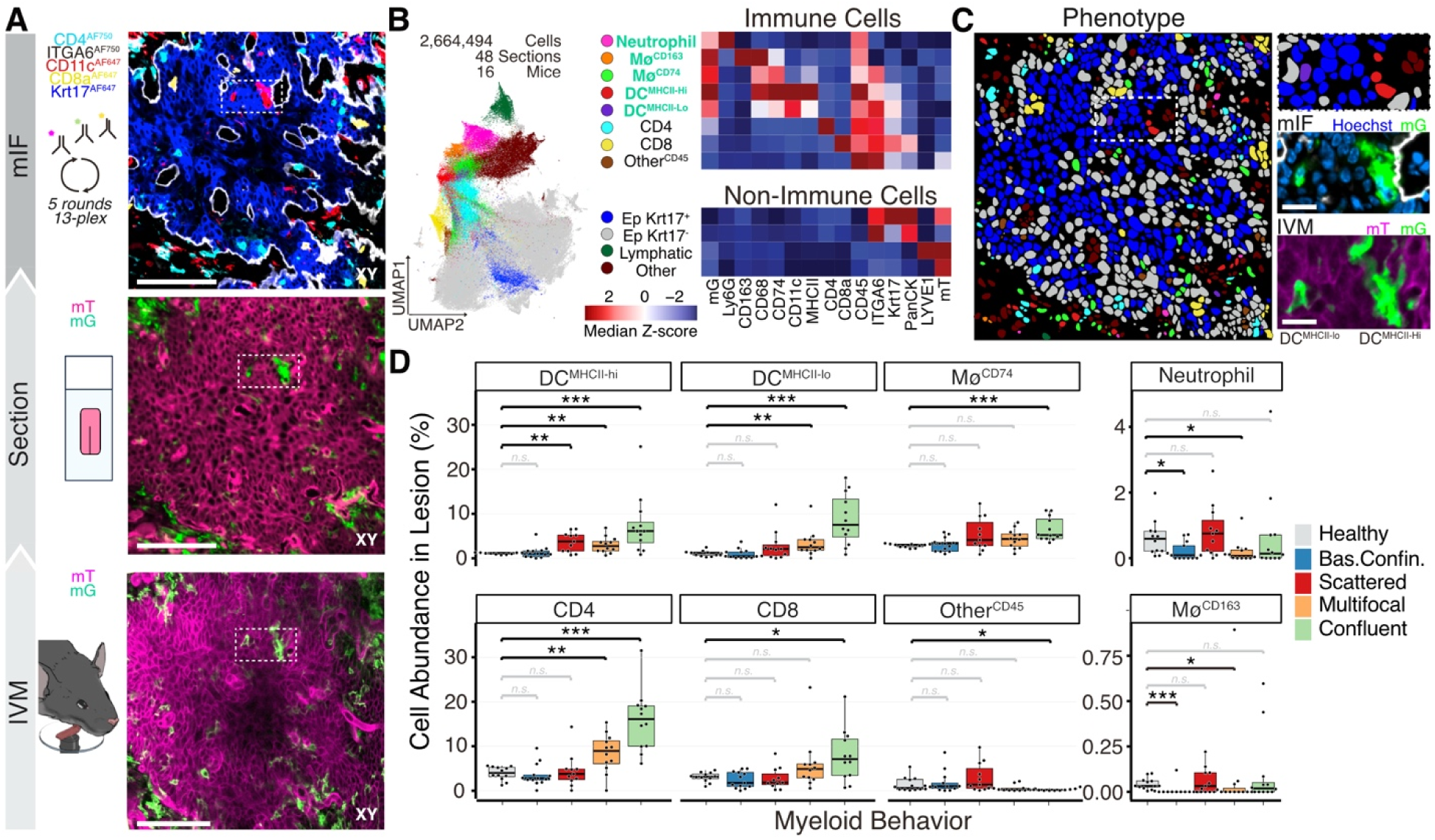
Correlative intravital microscopy and multiplex immunostaining identify tumor-infiltrating myeloid cells as antigen-presenting cells (APCs). Tongues from LysM-Cre; mT/mG mice at weeks 0, 8, 16, and 24 of the 4NQO regimen were imaged intravitally and immediately processed for cyclic multiplex immunofluorescence (mIF). **(A)** Schematic of the correlative workflow from intravital imaging to histological sectioning and mIF. **(B)** Unsupervised Phenograph clustering of single-cell mIF data displayed by UMAP, identifying 12 immune and non-immune cell phenotypes. Heatmap shows z-score normalized median expression level of each marker for each cell type. **(C)** Spatial correspondence of segmented cells between intravital and mIF datasets. Insets highlight the same myeloid cells identified by both modalities. Scale bar, 20 µm. **(D)** Quantification of immune cell subsets within 48 lesions stratified by intralesional myeloid organization. Pairwise Wilcoxon tests compare each organization to adjacent healthy basal epithelium in mice treated with 4NQO. mT/mG: Membrane-bound TdTomato/GFP.

Consistent with longitudinal intravital imaging, lesion size increased with treatment duration, whereas intralesional NADH intensity and myeloid infiltration declined over time (Fig. S4A–C). We identified a total of 55 lesions in the mIF datasets, of which 48 could be spatially matched to intravitally imaged lesions (Fig. S4D). Multiplex images were subjected to single-cell segmentation and unsupervised clustering using the *analysis of SPAtial single-Cell datasets* (SPAC) computational framework^38^ (Fig. 4B-C), resolving 12 major cellular phenotypes encompassing immune and non-immune compartments. Within the LysMcre^+^/GFP^+^ myeloid compartment, we identified neutrophils (Ly6G^+^), CD68^+^/CD74^+^ macrophages (Mø^CD74+^), CD68^+^/CD163^+^ macrophages, and two populations of dendritic cells distinguished by MHCII expression (CD11c^+/^MHCII^hi^; CD11c^+^/MHCII^low^). Both dendritic cell subsets exhibited strong LysM-driven GFP expression and elevated CD68 levels, indicating a myeloid origin consistent with monocyte-derived dendritic cells (mDCs). Additional immune populations included CD4⁺ and CD8⁺ T cells, as well as a residual CD45⁺ population lacking lineage markers. Analysis of non-immune compartments resolved healthy epithelium (ITGA6⁺/PanCK⁺/Krt17⁻), transformed epithelium (Krt17⁺), and lymphatic vessels (LYVE1⁺). B cells were rare or absent at all time points (data not shown), consistent with previous reports in both the 4NQO tongue model^39^ and human HNSCC^14^ and were therefore excluded from the final staining panel.

To link the immune composition with intravital myeloid organization, we quantified immune cell subsets in 48 lesions and stratified them by the dominant intralesional myeloid behavior observed by IVM (Fig. 4D). Mature and immature DCs, together with CD74⁺ macrophages, constituted the predominant infiltrating myeloid populations in lesions exhibiting confluent, multifocal, or scattered organization. In contrast, CD163^+^ macrophages and neutrophils were largely restricted to the stroma and were rarely detected within the epithelium compartment. Neutrophils were notably not enriched within lesions relative to healthy-adjacent epithelium, despite being abundant in the tongue upon 4NQO treatment (Fig. S4E-F).

Analysis of the lymphoid compartment revealed a distinct association with myeloid organization. CD4⁺ T cells were enriched in lesions containing multifocal or confluent myeloid clusters, whereas CD8⁺ T cells were selectively enriched in lesions exhibiting confluent organization, consistent with effective immune engagement. Lesions with scattered myeloid infiltration showed limited T-cell recruitment despite substantial myeloid presence.

Unsupervised clustering of immune compositions across all lesions defined six immune signatures that closely aligned with intravital myeloid behaviors (Fig. S4G-I). Signatures enriched in mDCs and CD4⁺ T cells predominated in lesions with confluent organization and were most frequent at earlier time points, whereas T cell–poor, mDC-dominated signatures increased over time and were associated with scattered organization. Immune-dormant signatures resembling healthy tissue remained stable across the treatment course (Fig. S4J).

Together, these data identify monocyte-derived dendritic cells and CD74⁺ macrophages as the principal myeloid populations infiltrating premalignant lesions. The spatial organization of these antigen-presenting cells, rather than their abundance alone, distinguishes immune-active lesions associated with regression from immune-poor lesions associated with progression.

### Distinct transcriptomic programs underlie infiltrating myeloid organization

Although both clustered and scattered infiltrates were dominated by antigen-presenting cells (APCs), these myeloid organizations differed in their capacity to recruit T cells. To determine whether these behavioral states could be distinguished transcriptionally, we performed GeoMx Digital Spatial Profiling (DSP) on endpoint samples collected at 24 weeks^40^ (Fig. 5A). Six sections spanning the epithelium and lamina propria were analyzed, including two lesions with scattered infiltrates and two with dense myeloid clusters that were spatially matched to intravital imaging data (Fig. 5B). Across 153 areas of illumination (AOIs), we profiled 19,962 genes covering lesions, interstitial tissue, and adjacent healthy epithelium. To account for the limited spatial resolution of DSP relative to single-cell RNA sequencing^40^, genes were assigned to myeloid (GFP⁺), epithelial (PanCK⁺), or other (GFP⁻/PanCK⁻) compartments based on their predominant expression. This classification identified 5,972 myeloid-associated genes, 11,548 epithelial genes, and 2,442 genes assigned to other compartments (Fig. S5A). As validation, tumor epithelium showed upregulation of Krt17 and downregulation of Aldh3a1 relative to healthy epithelium, confirming the neoplastic state^41,42^ (Fig. S5B). Tumor-derived myeloid AOIs were enriched for antigen-presentation genes such as Cd74, Ctss, and Cst3^43^ compared to interstitial myeloid cells (Fig. S5C), consistent with mIF identifying mDCs and CD74⁺ macrophages as the dominant infiltrating populations (Fig. 4D). Direct comparison of clustered and non-clustered tumor-derived myeloid AOIs revealed distinct transcriptional programs (Fig. 5C). Scattered myeloid cells from progressing lesions expressed high levels of Mmp12, ApoE, and LipA, signatures consistent with inflammatory remodeling, lipid-metabolic programs, and M2-like polarization^44,45^. In contrast, dense myeloid clusters from regressing lesions were dominated by interferon-driven programs, including Cxcl9, Cxcl10, and interferon-stimulated genes such as Isg15 and Irf7, known to mediate T-cell recruitment through CXCR3-dependent positive feedback with IFNγ^46^.

**Figure 5.**
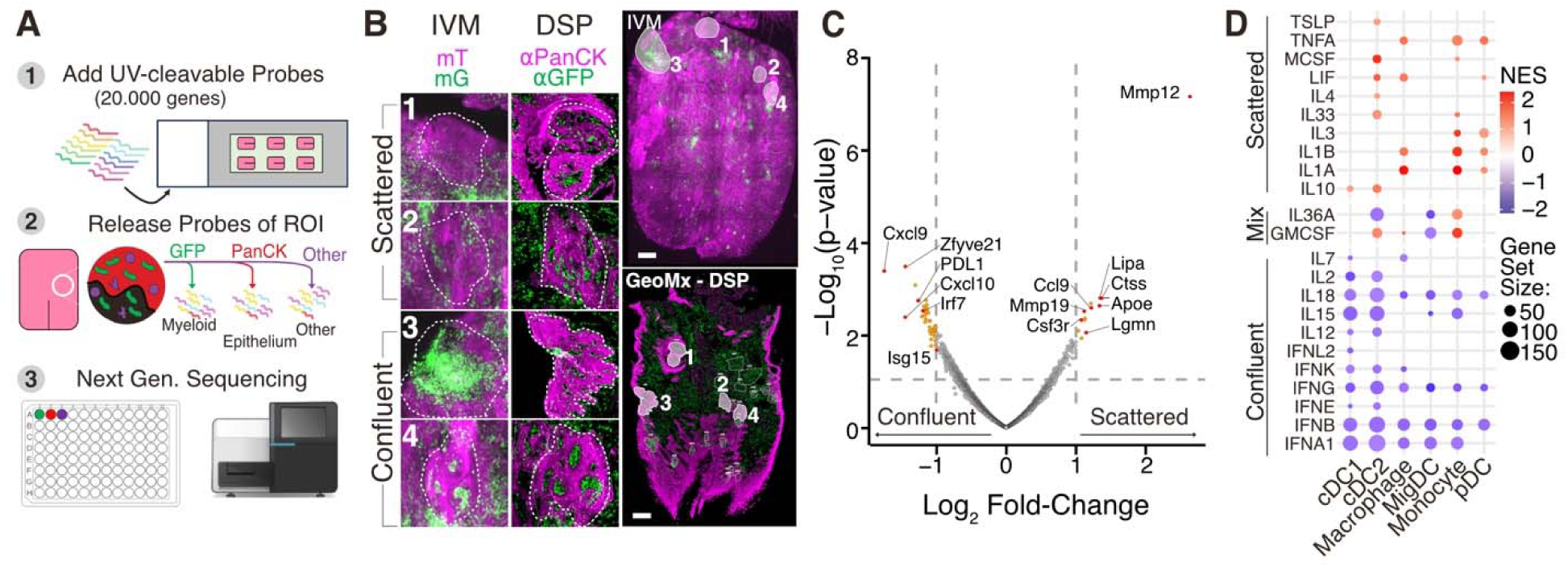
Correlative spatial transcriptomics reveals divergent transcriptional programs in clustered and scattered myeloid infiltrates. GeoMx Digital Spatial Profiling (DSP) was performed on a tongue harvested after 24 weeks of 4NQO treatment, as described in Fig. 4. (A) Schematic of the DSP workflow including whole-transcriptome profiling: (1) transcriptome-wide UV-cleavable RNA probes were hybridized to mRNA; (2) samples were stained for GFP and PanCK, and probes were collected from GFP⁺, PanCK⁺, or GFP⁻/PanCK⁻ (Other) regions, yielding 153 areas of illumination (AOIs); (3) collected probes were sequenced by next-generation sequencing (NGS). (B) Correlative intravital microscopy of a tongue containing four lesions after 24 weeks of 4NQO treatment. Insets show magnified images of clustering and non-clustering myeloid behaviors determined by IVM (left) and corresponding DSP sections (right) with enlarged views of the same regions. (C) Volcano plot showing differential gene expression between myeloid AOIs from clustered versus non-clustered lesions. A total of 5,972 myeloid-specific transcripts were analyzed (Fig. S5). Dashed lines indicate significance thresholds (|log₂FC| > 1, p < 0.05 by linear mixed model). (D) Gene Set Enrichment Analysis (GSEA) using the Dictionary of Immune Responses showing enriched profiles for monocyte, macrophage, and dendritic cell programs (FDR q < 0.25). mT/mG: Membrane-bound TdTomato/GFP.

Gene Set Enrichment Analysis (GSEA) using the Dictionary of Immune Responses^47^ confirmed distinct divergence (Fig. 5D). Scattered APCs were enriched for Interleukin 1-and tumor necrosis factor α-responsive gene signatures, whereas clustered myeloid cells from regressing lesions were enriched for interferon-driven, T cell–recruiting programs. Notably, PD-L1 expression was higher in confluent myeloid clusters than in scattered infiltrates, consistent with a regulated immune activation state rather than immune evasion^48^.

Together, these data show that spatially distinct APC organizations are associated with fundamentally different transcriptional states. Clustered APCs establish interferon- and chemokine-rich immune niches associated with lesion regression, whereas scattered APCs adopt lipid-metabolic and inflammatory programs associated with immune-poor, progressing lesions.

### Myeloid niches predict lesion onset

To determine whether myeloid cell behavior could predict lesion fate before morphological transformation, we examined myeloid behavior at early time points prior to detectable lesion formation. Whole-tongue intravital imaging revealed transient niches of myeloid cells that were not associated with visible lesions at any stage and were absent from untreated tongues (Fig. 6A–B). mIF analysis showed that these early niches were primarily composed of DCs and CD74⁺ macrophages, accompanied by CD4⁺ T cells (Fig. S6A–C), resembling the cellular composition of lesion-associated myeloid niches. Quantification of niche coverage across the ventral tongue revealed a rapid expansion phase during weeks 1-4, followed by a plateau between weeks 4 and 12 and a subsequent decline that initiated before withdrawal of 4NQO at week 16 (Fig. S6D). This decline likely reflects a reduced capacity to form myeloid niches rather than an acute response to carcinogen cessation. Spatial transcriptomic analysis of these orphan niches at week 24 revealed upregulation of *Cxcl9* and *Cxcl10* along proteins involved in antigen presentation, consistent with a T cell–recruiting phenotype (Fig. S6E).

**Figure 6.**
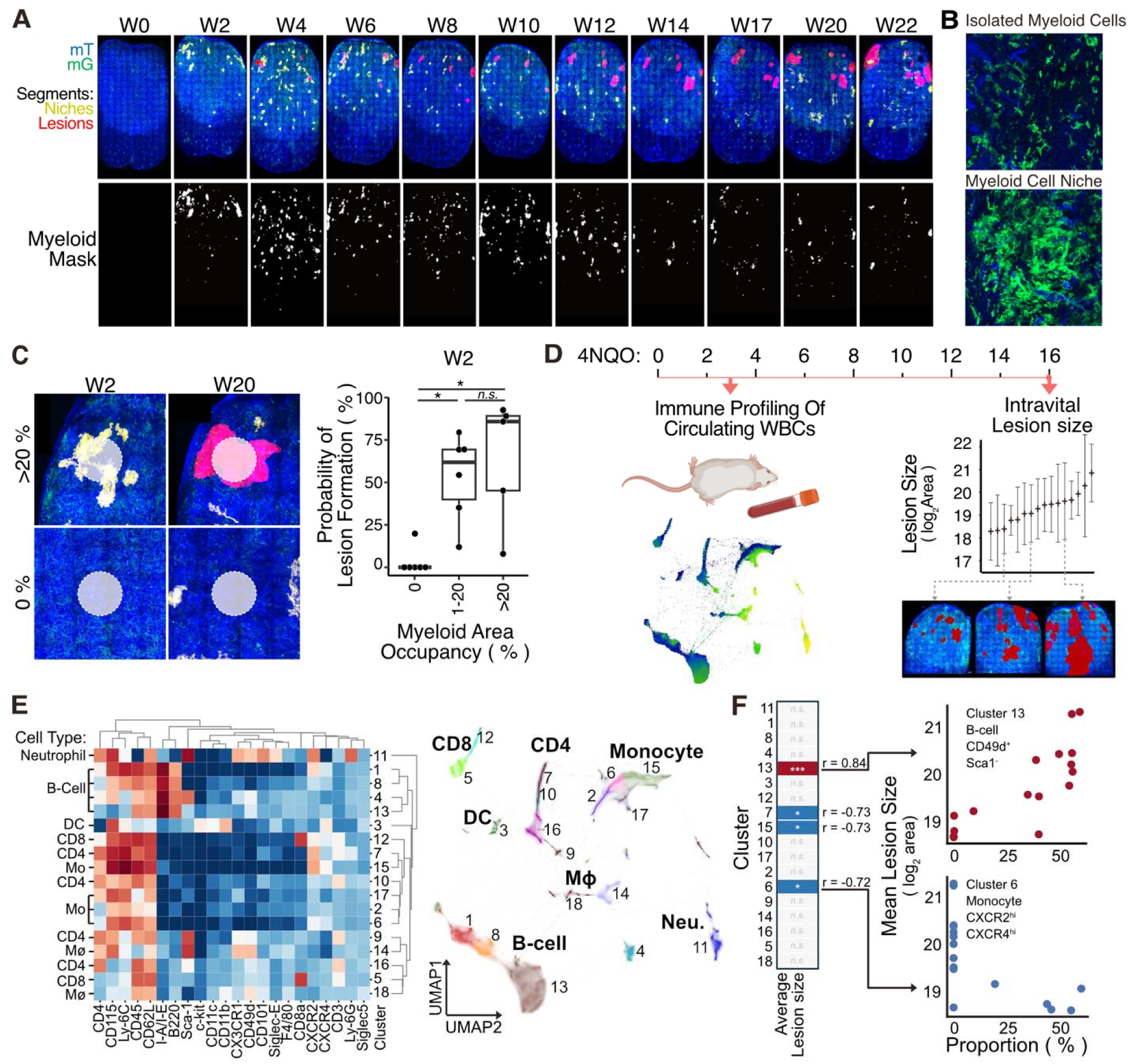
Myeloid niches predict lesion onset. (A) Quantification of myeloid niche coverage across the ventral tongue during 4NQO treatment, showing early accumulation (weeks 1–4), plateau (weeks 4–12), and decline thereafter. (B) High magnification images comparing niche-forming myeloid cells with isolated individual cells. (C) Probability of lesion formation as a function of myeloid niche density within a 250-µm radius at early time points. Box-plots showing probability of lesion formation grouped by no (0%), low (1-20%) or high (>20%) recruitment of myeloid niches at early timepoints. Datapoints represent individual mice. Pairwise Wilcoxon tests were performed to compare each group. **D)** Experimental design for peripheral blood profiling at week 3 and lesion assessment at week 16. Blood was collected from 16 mice at week 3 of the 4NQO treatment and subjected to multiplex flow-cytometry for immune phenotyping. The same mice underwent imaging at week 16 for lesion number and size. **(E)** Heatmap of marker expression after unsupervised clustering using the Hal-X algorithm. UMAP plot of their relative similarity is shown to the right. **(F)** Correlation between early circulating immune subsets and subsequent lesion burden. Two circulating immune subtypes are highlighted. High levels of circulating B-cell^CD49d+/Sca1+^ (Cluster 13) predict higher lesion burden at week 16 of treatment. High levels of Monocytes ^CXCR2+/CXCR4+^ predict low lesion burden at week 16 of treatment. r = Spearman correlation coefficient, * for p < 0.05, ** for p < 0.01, and *** for p < 0.001 after Bonferroni correction. mT/mG: Membrane-bound TdTomato/GFP.

Spatial mapping showed that niches frequently localized to sites that later developed pre-malignant lesions. Lesion onset probability correlated with the density of myeloid niches present within a 250 µm radius at early time points (week 2-3, Fig. 6C). Histological analysis further revealed that early myeloid clusters expressed elevated levels of PD-L1 and recruited FOXP3⁺ regulatory T cells (Fig. S6F-G), defining them as localized, immunoregulatory niches. These features suggest that early myeloid clustering reflects a transient, spatially confined immune response that may fail to eliminate premalignant epithelial cells and instead mark tissue regions permissive for lesion initiation.

To assess whether early local immune activity was reflected systemically, we profiled circulating white blood cells at week 3 of 4NQO treatment using multiplex flow cytometry and correlated the blood profiling results with lesion burdens at week 16 (Fig. 6D). Unsupervised hierarchical clustering analysis^49^ identified 18 circulating immune cell populations (Fig. 6E). A subset of monocytes (CXCR4+/CXCR2+) and CD4 T-Cells (CXCR2+/CD115+) showed statistically-significant negative correlations with subsequent lesion size, whereas the frequencies of a B-cell population (CD49d⁺/Sca1⁻) were strongly positively correlated with lesion burden (r_S_ = 0.84, p = 0.0008) (Fig. 6F), indicating that early systemic immune states of blood may partially reflect future lesion severity overall.

Together, these data show that transient myeloid clusters arise before detectable lesion formation and spatially precede the development of premalignant lesions. Their PD-L1–expressing, Treg-enriched nature suggests that early immune organization may create permissive niches for tumor initiation, providing both a spatial and temporal predictor of lesion onset.

## Discussion

This study demonstrates that the spatial organization of myeloid cells alone predicts the fate of premalignant lesions. By longitudinally following individual lesions in an immunocompetent 4NQO carcinogenesis model, we identified three distinct trajectories of lesion growth, progression, stability, and regression, each associated with a characteristic myeloid architecture. Correlative intravital imaging, multiplex immunostaining, and spatial transcriptomics revealed that lesion fate depends not on the mere presence of immune infiltrates but on their spatial and temporal organization within the epithelium.

Across all lesion trajectories, the dominant infiltrating myeloid populations consisted of monocyte-derived dendritic cells and CD74⁺ macrophages exhibiting antigen-presenting programs. However, these APCs adopted distinct organizational and transcriptional states. Dense, confluent APC clusters were characteristic of regressing lesions and associated with robust T-cell recruitment, interferon-responsive transcriptional programs, and expression of the chemokines CXCL9 and CXCL10. These features closely resemble the dendritic cell–T cell immune hubs described in human cancers, where their presence predicts a favorable response to immune checkpoint blockade^11,12,14^. Our findings extend this paradigm by showing that such immune niches are not only associated with therapeutic response but can also accompany spontaneous regression during early tumor evolution.

A key and unexpected finding was the presence of transient myeloid niches that formed weeks before detectable lesion onset. These niches were compositionally and transcriptionally similar to lesion-associated immune niches yet were enriched for PD-L1 expression and regulatory T cells. Spatial mapping showed that they preferentially localized to sites that later developed premalignant lesions. These observations suggest that early myeloid niches reflect abortive or incomplete immune surveillance, creating spatially confined sites that are permissive for tumor initiation when epithelial cells escape clearance. The dual presence of immune activation signals and immune regulatory mechanisms within these clusters highlights a fragile equilibrium between rejection and tolerance at the earliest stages of carcinogenesis.

These findings have important implications for head and neck squamous cell carcinoma, a disease with a high global burden and limited predictors of progression from premalignant lesions^50,51^. While immune checkpoint blockade has transformed the treatment of advanced HNSCC, only a minority of patients respond durably^52^. Our data suggest that immune architecture, rather than immune cell abundance, may determine whether early lesions are cleared or progress, providing a potential explanation for heterogeneous responses to immunotherapy. Importantly, these immune architectural changes may be reflected in peripheral blood immune cell states, potentially enabling early prediction of lesion progression. Moreover, the identification of early APC states and transient immune niches raises the possibility that tumor fate may be predicted, and potentially altered, before malignant transformation occurs.

From a translational perspective, our results suggest several opportunities. First, detection of CXCL9/10⁺, PD-L1⁺ myeloid clusters or MMP12-expressing APC states could identify high-risk fields predisposed to lesion development. Second, strategies that stabilize productive immune niches or prevent early APC dysregulation may enhance spontaneous regression and improve responsiveness to immune checkpoint blockade. Finally, advances in endoscopic multiphoton microscopy and metabolic imaging may enable longitudinal monitoring of immune architecture and epithelial metabolism in patients^53,54^, translating the principles identified here into early detection and cancer interception strategies.

In summary, this work establishes spatiotemporal myeloid organization as a fundamental determinant of premalignant lesion fate. By resolving immune dynamics before, during, and after lesion emergence, we provide a mechanistic framework for spontaneous regression and progression and identify early immune states that may be exploited for cancer prediction and prevention.

## Materials and Methods

### Animals and procedures

All experiments were approved by the National Cancer Institute (National Institutes of Health, Bethesda, MD, USA) Animal Care and Use Committee and were compliant with all relevant ethical regulations regarding animal research. LysMcre (B6.129P2-Lyz2tm1(cre)Ifo/J) and mT/mG (B6.129(Cg)-Gt(ROSA)26Sortm4(ACTB-tdTomato,-EGFP)Luo/J were purchased from Jackson Laboratory and backcrossed at least 6 generations into c57Bl6/NJ background before crossed to homozygosity for both gene elements. All mice used in this study were between 8 and 14 weeks of age at the start of treatment. Mice were anesthetized by isoflurane inhalation (3%) followed by an initial intraperitoneal injection of a mixture of ketamine (100lmg per kg) and xylazine (20lmg per kg) in lactated Ringer’s solution before imaging sessions and maintained as necessary during imaging.

### 4NQO chemical induction of tumors

Premalignant lesions and tumors were induced as previously described^29^. In brief, 4-Nitroquinoline 1-oxide was administered at a concentration of 50µg/mL in the drinking water for 16 weeks (week 1-16) followed by a recovery period with regular water for 8 weeks (week 17-24). To maintain weight, the mouse diet was supplemented with a soft transgenic dough-diet (Bio-serv, S3472) once a week starting at week 5 of treatment. An equal number of male and female mice were used.

### Intravital Microscopy

Two-photon microscopy was performed using an inverted laser-scanning two-photon microscope (MPE-RS; Olympus, Center Valley, PA) equipped with a tunable laser (Insight DSþ; Spectra Physics, Santa Clara, CA). The tongue of the anesthetized animal was gently placed in a 3D-printed tongue holder device (Wang et al. unpublished) with the ventral side facing the objective. The entire ventral area was imaged using an Olympus UPlanSApo IR 30x silicone oil objective (NA: 1.05, Evident) or a UPLXAPO 10x air objective (NA: 0.4, Evident) using a resonance scanner set to 3-line repetitions. The most superficial 150µm was acquired in each stack at a 2µm distance between optical slices (z-resolution). Excitation was performed at 900 and 740 nm, and the emitted light was collected by an appropriate set of mirrors and filters on three detectors (bandpass filters: blue for collagen or NADH: Z 410 to 460 nm; green for GFP: Z 495 to 540 nm; and red for TdTomato: Z 575 to 645 nm). The resulting stacks were stitched using the Fluoview software (Evident) after each imaging session. The same imaging setup was used for time-lapse imaging, except a 1.5x digital zoom and 6 line repetitions were used for an imaging depth of 72µm (49 optical sections at a z-resolution of 1.5µm per optical slice). 5-10 min of timelapse was acquired at a framerate of 19.4 3D stacks per second. High-resolution images were obtained in Galvano scanning mode using a UPLANSAPO 40X silicon immersion objective (NA: 1.40, Evident).

### Intravital Analysis

Intralesional measurements - Stitched whole tongue 3D images were analyzed for lesion size using the Imaris v9.2.1 software (Oxford Instruments). First, the spatial locations of lesions were identified by three criteria: 1) altered epithelial basal cell morphology, 2) degree of collagen displacement compared to surrounding epithelium, and 3) maintained dysregulated morphology and/or displaced collagen for at least 3 timepoints in the same spatial location. To segment lesion volumes, the epithelial compartment was masked using the collagen channel as a reference, followed by manual outlining of the lesion. The collagen channel was first segmented using the surfaces tool setting smoothing to 10µm and local threshold to 2, selecting the largest surface from the results. The TdTomato channel was masked by the negative collagen surface, producing a channel with isolated epithelium. Each lesion was manually segmented in a 2-step process. First, the outline of the transformed epithelium in the masked TdTomato channel was manually annotated and masked using the surfaces tool; to generate a volumetric mask that follows the contour of the surface of the epithelium, the resulting lesion channel underwent an additional automated surface detection without smoothing and thresholds were set to 120 for S3 and S4 tongues, and 90 for S1 and SW tongues. The volume of the resulting surface was exported and analyzed using R. Lesion volumes were log2 transformed and fit to a linear equation over time in weeks using the lm() function (y= bx + a) and lesions with >0.5 *r*^2^ and *b* > 0.1 were characterized as progressing. Lesions were characterized as regressing when endpoint volume was ≤ 25% of the peak volume, and stable in cases where the fit was <0.5 *r*^2^ and *b* < 0.1, while endpoint size was more than 25% of the peak volume.

To measure Myeloid displacement for each lesion, the GFP channel was masked using the established lesion segments using Imaris. Myeloid cells were segmented using the surfaces tool with smoothing set to 3 and local threshold set to 8, followed by a voxel filter of >=10. The resulting myeloid segment parameters within each were exported, and displacement was calculated as the total myeloid segment volume divided by the whole lesion volume.

Myeloid behavior was determined from manual assessment of the 3D volume of each lesion time point using Imaris. The lesions were grouped as follows. “Basally confined”: All Myeloid cells within the lesion epithelium touched the basement membrane, and myeloid cells were neither touching each other nor found above the parabasal layer. “Scattered”: Myeloid cells were found in the spinous layer, scattered evenly throughout the epithelium without touching each other. “Multifocal”: One or more small clusters of myeloid cells are touching cell body-to-cell body. “Confluent”: A central, multicellular structure made out of myeloid cells.

Intralesional NADH level was determined from qualitative assessment of the whole-tongue scans acquired at 740nm according to the following criteria. “Low”: at least one coherent spot within the lesion is low in NADH signal in the entire column from corneal to basal layer. “Normal”: NADH level in the lesion was at the same level as adjacent healthy tissue. “High”: NADH level in the lesion is visibly more intense than in adjacent healthy tissue. “Slightly down” No coherent spot within the lesion had visibly lower NADH from corneal to basal layer, but the NADH level does appear lower than healthy adjacent tissue.

Myeloid recruitment - To compare myeloid recruitment in tongue images between timepoints, each scanned image was imported using BioFormats, ImageJ (NIH) at the resolution level 2, and the 3D volumes were converted into sum-projects. To generate the myeloid niche mask for each sum-projection, the GFP-channel was smoothened at a 10µm radius using the Gaussian algorithm, thresholded using the triangle method, and the papillae area was manually annotated and excluded. The same anatomical landmarks (Blood vessels and muscles) were manually labelled at each time point. The image registration of every two consecutive time points was accomplished by matching these landmarks to achieve the minimum mean squared errors, assuming that there are only translational motions, rotations, and stretching or compressing along the orthogonal directions of the same tongue between the consecutive images. The optimization process of the image registration was implemented with a customized software coded in Python with the Pytorch module. The final mean errors to register the anatomical landmarks were less than 40 µm between two consecutive time points.

By combining the lesion masks and the image registration results, for certain time point t, we defined the healthy mask of the papillae-less area according to the following criteria: 1) pixels in the mask appear in every time point after time point t (including t); 2) pixels in the mask are all at least 250 µm away from the boundary of the lesion mask or the manually tracked locations of lesions.

For each pixel in the 2D z-projection image, a local circular region (ROI) of interest with a 250 µm radius was drawn around it. The fraction of the area of this ROI occupied by the myeloid niches was calculated from the 2D z-projection masks of the myeloid clusters. Thus, the distribution of the fraction of peripheral ROI occupied by the myeloid cells over the entire tongue was generated at each time point.

For all pixels at time point t, we considered the probabilities of the two following conditions: A) The location at the tongue corresponding to the pixel in the image is healthy in later time points (including t) during the longitudinal experiments. B) The local circular ROI with 250 µm around the occupied by the myeloid niches by a fraction of at least x%. The probability of condition A, P(A), can be estimated by calculating the ratio of the healthy mask over the total mask of the papillae-less area imaged in every later time point (including time point t). The probability of condition B, P(B), can be estimated with the distribution of the fraction of the peripheral ROI occupied by the myeloid cells within the papillae-less area. By investigating the distribution of the fraction of peripheral ROI occupied by the myeloid cells within only the healthy mask, the conditional probability P(B|A) can be calculated according to Bayes’ theorem, P(A|B) = P(B|A)P(A)/P(B). Therefore, we can decide the probability of lesion onset after time point t (including t) if the local circular ROI with 250 µm around the occupied by the myeloid clusters by a fraction of at least x% as 1-P(A|B).

### Cardiac fixation and histological sectioning of the tongue

For cardiac perfusion, the left ventricle of the heart was punctured, and PBS supplemented with heparin (10U/mL, Hospira NDC0409-2720-30) was perfused to wash out the blood from the right atrium, followed by 10lml fixative (4% paraformaldehyde in 200mM HEPES, pH 7.3). After a total of 10 min of fixation, the animal was further perfused with 10mL quenching buffer (20mM glycine, 50mM Ammonium chloride in PBS), 10mL sucrose (15% in PBS), and finally 10 mL sucrose (30% in PBS). The tongues were excised and incubated in 30% sucrose on ice until the tissue sank in the solution. The tongue was snap-frozen in a modified embedding medium for frozen sectioning^55^ (7.5% HPMC 40-60cP, sigma H8384 & 2.5% PVP Mn m/w 360kDa, sigma PVP360, dissolved in water) on powdered dry-ice between two histology slides to maintain a flat orientation of the ventral epithelium. The glass slides were removed by gentle heating with a thumb on the ventral side, and 5µm cryosections were cut (Leica CM1860 cryostat) and collected on charged microscopy slides (Globe, #1358).

### Multiplex Immunofluorescence staining and imaging

Slides retrieved from −80C were equilibrated at −20C for 10min followed by 10 min at RT. Sections were post-fixed in 4% PFA (200mM HEPES pH 7.3) for 10 min at RT and washed 3 x 5 min (2x in 20 mM Glycine, 50mM NH_4_Cl in PBS and 1 x in PBS). Blocking was performed in 3% BSA, 5µg/mL Fc-Block (anti-CD15/32 TruStain FcX, Biolegend) for 1h at RT. For multiplex staining, the blocked sections were subjected to 6 rounds of the following procedure using different antibodies in each round (Round 1: CD45-AF75, CD8-AF647, Round 2: CD4-AF750, CD11c-AF647, Round 3: CD68-AF750, Ly6G-AF647, Round 4: LYVE1-Dy755, CD74-AF647, rabbit-anti Krt17-non conjugated. Round 5-1: Anti-rabbit-AF647, PanCK-AF555, ITGA6-Bio. Round5-2: Strep-AF750, MHC-II-AF594. Round 6: CD163-PE). For each round, slides were incubated with the antibodies diluted in 1% BSA, 1.25µg/mL Fc-Block for 2h, followed by 3x wash steps in PBS. Nuclei were stained using 5µg/mL Hoechst 33342 (Thermo Fischer Scientific) in PBS for 10 min followed by 2x wash steps in PBS and 2x wash steps in deionized water. Slides were mounted in Slowfade Diamond media (S36963, Thermo Fischer Scientific) and imaged using a Zeiss Axioscan Z.1 Slide Scanner equipped with a 20x objective (Plan-Apochromat, NA 0.8) and a Colibri 7 Flexible Light Source. After imaging, the slides were retrieved, and the coverslip was removed by submerging in PBS for 2 h. Fluorophores were bleached using 1mg/mL LiBH4 in deionized water for 15 min, followed by 3 wash steps in PBS before initiating the following round of staining and imaging. During round 5, an additional bleaching step for TdTomato was included by incubating in 3.5% H2O2 in 90% MeOH on ice for 20 min after the LiBH_4_-bleaching and washing steps. Additionally, the antibody incubation step (Round 5-1) was followed by 3 x wash steps in PBS and an additional secondary antibody incubation step (Round 5-2) for 2h at RT.

Hematoxylin and eosin stainings were performed by VitroVivo.

### Image analysis of multiplexed images

#### Image Preprocessing

Zeiss ZenBlue software was used to split each slide scan into individual tissues, and the czi files were loaded into the HALO v4.1 software (Indica Labs) for sequential stained image fusing and cell segmentation. Each round of tissue staining was merged by using the “register images” function using the Hoechst staining of the 6 sequential images for registration, followed by “fuse serial stain” with the Hoechst channel as reference and the first round as the image registration target. The resulting fused images were used for all downstream analysis. The outline of the ventral epithelial area and each lesion identified by IVM were manually annotated, and lesion annotations were subtracted from the intravitally imaged area to generate a “Healthy adjacent tissue” annotation.

#### Single Cell Segmentation

A classifier pipeline containing four individual dense-net v2 AI-based classifiers was used to segment in-focus and correctly merged extracellular matrix and epithelial tissue areas. The first classifier to determine tissue from glass was trained on 5 annotations of tissue and glass from separate tissues using the TdTomato channels of the first round, and segments containing tissue were selected. The second classifier, designed to recognize in-focus vs out-of-focus tissue, was trained on 12 areas in focus and 28 out-of-focus areas from 12 tissues, and the in-focus segments were selected. The third classifier for detecting correctly merged vs incorrectly merged areas was trained on 12 correctly merged areas and 17 incorrectly merged areas from 3 sections for the training using the Hoechst channels from the 6 images, and the correctly merged segments were selected. The fourth classifier for classifying tissues into ECM, Epithelium, and Muscle segments was trained on 9 ECM areas, 13 muscle areas and 109 epithelium areas from 9 tissues, and both ECM and epithelium segments were selected the final analysis. To segment nuclei within each segment, a Nuclei Seg (Plugin) - FL v1.0.0 classifier was trained on 446 nuclei from 5 areas sampled from 5 different tissues. To extract single-cell fluorescence intensities of 24 markers (including 6 rounds of Hoechst channels and 4 rounds of TdTomato channels) a HighPlex FL v4.3.2 module was used on all annotations. This module implemented the trained classifier pipeline to identify tissue regions and the nuclear segmentation plugin to identify cells using a fixed maximal 2 µm cytoplasmic radius for each cell. Whole-cell intensities were exported and used for downstream analysis.

#### Single cell analysis

Exported single cell values underwent unsupervised clustering for cell identity determination using phenograph as part of the “analysis of SPAtial single-Cell datasets” (SPAC) environment within the NIH Integrated Data Analysis Platform ^38^. First, batch correction was performed by applying arcsinh and z-score normalization. The normalized data was clustered using the phenograph node and transformed into the UMAP node in parallel. The resulting clusters were evaluated based on expression level, and all clusters with high expression of GFP, CD11c, CD68 and MHCII were pooled and underwent a second phenograph clustering to separate macrophages, dendritic cells, and misclassified epithelial cells. The percentage of immune cells within tumors was calculated using the epithelial segment of the manual annotations, and the results from different sections of the same lesion were summed before calculating the cell-specific prevalence.

### GeoMx Digital Spatial Profiling

Six sections from the same tongue of a female mouse that underwent the complete 24 weeks 4NQO-treatment regimen were collected with 25µm spacing between each section. For the NanoString GeoMx DSP RNA assays, slides were prepared following the Leica Biosystems BOND RX FFPE RNA Slide Preparation Protocol described in the GeoMx NGS Slide Preparation User Manual (NanoString, MAN-10L115-04). Briefly, slides were baked overnight at 60°C and then loaded into the Leica BOND RX device. Slides were treated sequentially following Leica BOND RX default HIER (ER2, 20 min, 95°C) Protocol and GeoMx DSP RNA Slide Prep Protocol (1 mg/mL proteinase K (Ambion, cat. 2546) in 1X phosphate-buffered saline (PBS) at 37°C for 15 min). After pretreatment, the slides were hybridized with the GeoMx Mouse Whole Transcriptome Atlas Mouse RNA for Illumina Systems (Nanostring, GMX-RNA-NGSMsWTA-4). The slides were dried of excess 1X PBS, set in a hybridization chamber lined with Kimwipes wetted with Diethyl pyrocarbonate(DEPC)-treated water, and covered with 200 μL prepared Probe Hybridization solution. HybriSlips (Grace Biolabs, cat. 714022) was gently applied to the slide, and incubated at 37°C overnight.

After hybridization, the HybriSlips were removed by dipping the slides in 2X saline-sodium citrate (SSC) (Sigma-Aldrich, cat. S6639)/0.1% Tween-20. To remove unbound probes, the slides were washed twice in Stringent Wash (50% formamide (ThermoFisher, cat. AM9342)/2X SSC) at 37°C for 25 min, followed by two washes in 2X SSC for 2 min. Slides were blocked in 200 μL Buffer W (NanoString), placed in a humidity chamber and incubated at room temperature for 30 min. The slide was dried of excess Buffer W, set in a humidity chamber, covered with morphology marker solution containing anti-GFP antibody (Abcam, Ab6673) and left to incubate at room temperature for 1 hour staining followed by 2x wash steps with SSC for 5 min. Secondary anti-goat antibody-594 and anti-pan-cytokeratin Alexa 532 antibody for 1 hour staining followed by 2x wash steps with SSC for 5 min. The sections were immediately loaded into the GeoMx instrument after staining.

### Digital Spatial Profiling analysis

Briefly, RNA probe-hybridized and antibody-stained slides were scanned with a 20X objective, collecting data using FITC/525 nm (excitation 480/28 nm, emission 516/23 nm), Cy3/568 nm (excitation 538/19 nm, emission 564/15 nm), Texas Red/615 nm (excitation 588/19 nm, emission 623/30 nm), and Cy5/666 nm (excitation 645/19 nm, emission 683/30 nm) channels. Regions of interest (ROI) were selected from the corresponding intravital images based on morphology as well as cell surface marker staining. Within each ROI, areas of illumination (AOI) were identified for RNA collection for either Myeloid cells (GFP+), Epithelial cells (PanCK+), or “other” cells (PanCK-, GFP-). ROIs were annotated directly in the DSP software, and UV light was used to cleave the barcode linkers of the prebound RNA probes in each AOI, and the cleaved probes in DSP were retrieved into collection plates.

DSP collection plates were frozen and stored at −80°C. Plates were thawed at room temperature and libraries were prepared per manufacturer’s guidelines. Collection plates were sealed with a semi-permeable membrane and dried down at 65°C for 1 hour. Each well was resuspended in 10 µL DEPC-treated water, incubated at room temperature for 10 min, and then spun down. The library preparation was carried out in a 96-well PCR plates by mixing 2 μL PCR mix (NanoString), 4 μL of index primer mix (NanoString), and 4 μL of DSP sample. The following PCR program was used to amplify the Illumina sequencing compatible libraries: 37°C for 30 min, 50°C for 10 min, 95°C for 3 min, followed by 18 cycles of (95°C for 15 s, 65°C for 1 min, 68°C for 30 s), 68°C for 5 min and a final hold at 4°C. A total of 6 plates of 96 wells each were used.

The indexed libraries were pooled with an 8-channel pipette by combining 2 μL per well from the 12 columns (one 96-well plate) into 8-well strip tubes and then pooled into a 1.5 m tube. The combined 50 µL pools were incubated with 60 μL SPRIselect beads (Beckman Coulter, cat. B23318) (1.2X bead to sample ratio) for 5 min in a 1.5 mL tube followed by standing on a magnetic stand for 5 Lmin before removal of the supernatant. The beads were then washed twice with 200 µL of 80% ethanol and air dried for 3 min before being eluted with 50 µL elution buffer (10 mM Tris-HCl pH 8, 0.05% Tween-20, Teknova cat. T1485). Then a second round of SPRIselect beads (Beckman Coulter, cat. B2331860 ul; 1.2X bead to sample ratio) selection was carried out directly on the bead suspension as above. The washed beads were eluted in 20 µL elution buffer (10 mM Tris-HCl pH 8, 0.05% Tween-20, Teknova cat. T1485), and 18 µL of the supernatant was extracted to a new tube. The 18 µL of clean-up supernatant from each 96-well plate was pooled, and the library fragment size was assessed with the D1000 Tape Station assay (Agilent Technologies) and the expected size of ∼162 bp was observed.

Total target counts per DSP collection plate for sequencing were calculated based on the NanoString DSP Worksheet. The target sequencing depth was 100 counts/um2. Libraries were sequenced on the NextSeq 2000 instruments with pair end 27×8×8×27 (CCR Genomics Core, Bethesda, Maryland, USA), and fastqs for each AOI were converted to Digital Count Conversion files (DCC files) using NanoString’s GeoMx NGS Pipeline v.2.3.3.10.

DSP analysis was performed using the R Package DSPWorkflow (https://github.com/NIDAP-Community/DSPWorkflow). All DCC files were combined with annotations and probe IDs mapped to gene names for the Mouse WTA probe set version 1.0 using the Nanostring R package GeoMxTools^56^. QC checks were applied in R studio using Nanostring’s guidelines for best practices of analysis of DSP data from RNA probes with NGS sequencing (GeoMx DSP Data Analysis User Manual (v3.1.2 software). 2024.). Upper quartile normalization was applied to all raw read counts after QC. PCA plots were used to evaluate the effect of normalization and to check for batch effects using the PCAtools R package^57^. 17 AOIs were removed to avoid technical noise after initial PCA indicated clustering for these AOIs from low nuclei and gene detection. Differential expression analysis was performed using GeoMxTools’s mixed model function with the slide number used as a random intercept, and a random slope was added when comparing AOIs within the same slide^58^.

### Analysis by spectral flow cytometry (CyTEK)

One to two drops of blood from the mouse cheek were collected into a blood collection tube. (Sarstedt EDTA). Ice-cold FACS buffer (4% of FCS and 0.1% of Sodium Azide in PBS) was added to the tube to transfer the blood samples to a 96-well plate on ice. After a brief centrifugation at 200 × g at 4°C for 1 min, the supernatant was discarded and cells were resuspended in 100 µL of ice-cold ACK (Ammonium/Chloride/Potassium) lysis buffer (Quality Biologicals, Gaithersburg MD), then incubated for 1 min on ice, followed by washing. The blood cells were then resuspended in ice-cold antibody mix (Table S1) and incubated for 30 min on ice. After washing, cells were resuspended in 100 µL of ice-cold 1.6% PFA buffer and incubated for 15 min. Following a final wash, cells were resuspended in 100 µL of FACS buffer and analyzed by CyTEK. Raw .fcs files were first normalized with reference single-cell staining controls and then gated on singlet cells for analysis. Channel values from single cells were exported from CyTEK and subjected to clustering analysis. To characterize cell types, we applied HAL-x^49^, a custom, scalable, hierarchical clustering algorithm. A key advantage of HAL-x is its ability to train a clustering model on a subset of cells (e.g., 10) and then rapidly apply the trained model to cluster the entire dataset (>10 cells) and/or new datasets.

## Acknowledgments

This research was supported [in part] by the Intramural Research Program of the National Institutes of Health (NIH). The contributions of the NIH author(s) were made as part of their official duties as NIH federal employees, are in compliance with agency policy requirements, and are considered Works of the United States Government. This project has been funded in whole or in part with Federal funds from the National Cancer Institute, National Institutes of Health, Department of Health and Human Services, under Contract No. 75N91019D00024. However, the findings and conclusions presented in this paper are those of the author(s) and do not necessarily reflect the views of the NIH or the U.S. Department of Health and Human Services, nor does mention of trade names, commercial products, or organizations imply endorsement by the U.S. Government. T.D.M is supported by a Novo Nordisk Foundation Grant NNF0067602. We thank the NCI Center for Cancer Research (CCR) Spatial Imaging Technology Resource (SpITR) and Laboratory Cancer Biology and Genetics (LCGB) Microscopy Core for technical support as well as the CCR Genomics Core, for performing RNA-seq. We also thank Dr. Amiran K. Dzutsev at the Laboratory of Integrative Cancer Immunology, NCI, NIH, for critical reading and Dr. George Zaki at the Biomedical and Computational Science Directorate, Frederick National Laboratory for Cancer Research, for assistance with analysis performed in SPAC.

## Author contributions

RW and TDM for Conceptualization. TDM, DC, MOH, Emily Chen, RL, NK, WW, DJ, GAB, RW, for Methodology, TDM, DJ, DC, MOH, SMH, MW, DT, MH, Emily Chen, SAE, NK, WW, DG, and GAB for investigation. TDM, DC, NK, RW for imaging. TDM, RW for funding acquisition. TDM, RW for Project administration. TDM, RW wrote the original draft. All authors edited and agreed to the final version of the manuscript.

## Additional Information

Correspondence and requests for materials should be addressed to Roberto Weigert.

