## Supplementary Figures for "Myeloid States Predict Fate of Premalignant Lesions"

### Longitudinal Imaging of the Premalignant Tumor Microenvironment Reveals Transient Myeloid States Predictive of Tumor Fate

Thomas D. Madsen[1,2*], Dongya Jia[3], Desu Chen[1], Maria O. Hernandez[4], Sarah M. Hammoudeh[1], Edmund Cauley[5], Marco Heydecker[1], Desiree Tillo[6], Madeline Wong[6], Emily Chen[1], Salma Abu-Elnaj[1], Ross Lake[7], Noemi Kedei[4], Gregoire Altan-Bonnet[3] Weiye Wang[1], Roberto Weigert[1*]

1. Laboratory of Cellular and Molecular Biology, Center for Cancer Research, National Cancer Institute, National Institutes of Health, Bethesda, MD, USA
2. Copenhagen Center for Glycomics, University of Copenhagen, Department for Cellular and Molecular Medicine, Copenhagen, Denmark
3. Immunodynamics Group, Laboratory of Integrative Cancer Immunology, Center for Cancer Research, National Cancer Institute, Bethesda, MD, USA.
4. Spatial Imaging Technology Resource, Center for Cancer Research, National Cancer Institute, National Institutes of Health, Bethesda, MD, USA
5. Advanced Biomedical Computational Sciences, Frederick National Laboratory for Cancer Research, Frederick, MD, 21702, USA.
6. CCR Genomics Core, Center for Cancer Research, National Cancer Institute, National Institutes of Health, Bethesda, MD 20892, USA.
7. Laboratory of Cancer Biology and Genetics, Center for Cancer Research, National Cancer Institute, National Institutes of Health, Bethesda, MD, USA

*Corresponding authors:

**The PDF file includes:**

Figure S1-S6

Legends for Video S1-S2

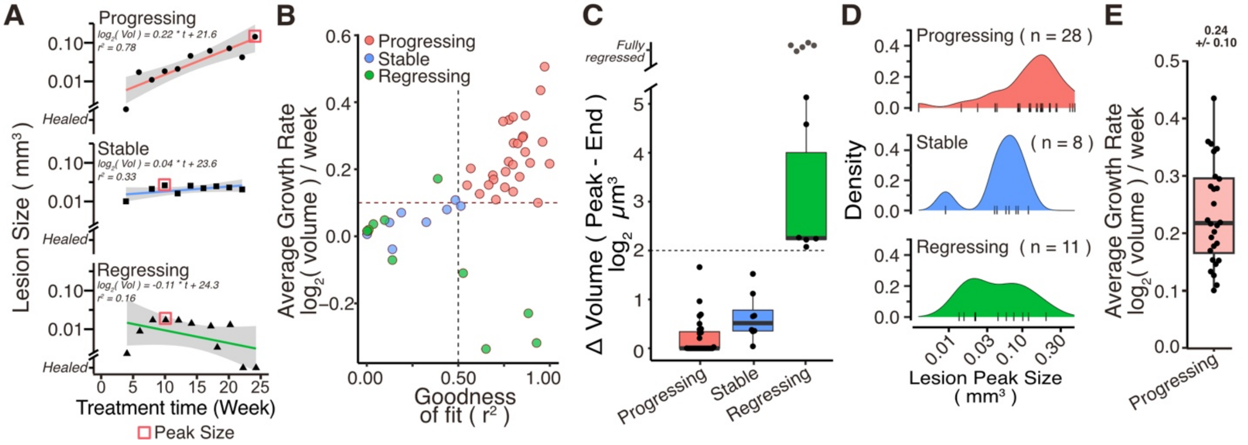


**Supplementary Figure 1 – Volumetric analysis of premalignant lesion growth characteristics and myeloid infiltration.** Tongues from mice treated with 4NQO were imaged by multiphoton microscopy, and lesion volumes were quantified as described in Figure 1. Lesion trajectories were classified as progressing, regressing, or stable by fitting lesion volumes to an exponential growth curve. **(A)** Representative linear fits of $\log_2$-transformed lesion volumes illustrating the three trajectories. Progressing lesions show a fit with $r^2 > 0.5$, whereas regressing and stable lesions show $r^2 < 0.5$. The peak size indicates the time point at which the lesion reached its maximal volume within the duration of the experiment. **(B)** Summary of the goodness of fit ($r^2$) and slopes of exponential growth curves for all lesions from eight mice. **(C)** Boxplots showing the difference between the endpoint volume and the maximum (peak) volume for each lesion. Lesions that shrink to ≤ 25 % (dashed line) of their peak volume are classified as regressing. **(D)** Density plot showing the distribution of peak lesion volumes as defined in (A). Regressing and stable lesions have comparable peak sizes, whereas progressing lesions are larger. **(E)** Boxplot of growth rates for progressing lesions, with an average volumetric increase of 24 ± 10 % per week across all progressing lesions.

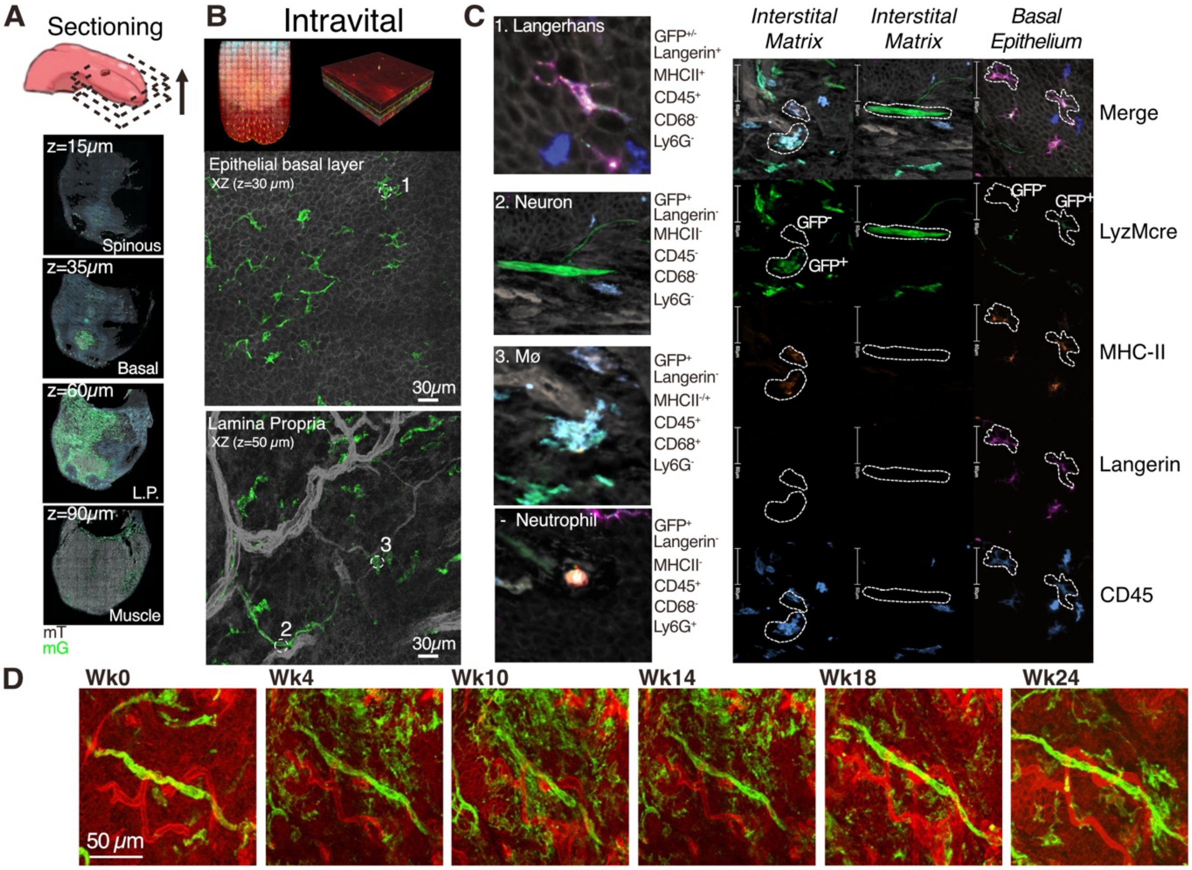


**Supplementary Figure 2 – Characterization of immune cells in the healthy tongue. (A)** Ventral sectioning of the tongue enables staining of all epithelial layers while preserving the same orientation used for intravital microscopy imaging. mT: TdTomato, mG: GFP **(B)** Intravital images showing GFP$^+$ myeloid cells within the epithelial layer and the interstitial matrix. **(C)** Immunofluorescence analysis of the epithelium and interstitial matrix identifies three major GFP$^+$ myeloid populations: (1) Langerhans dendritic cells residing in the epithelium, (2) neurons, and (3) monocytic dendritic cells located within the interstitial matrix. Neutrophils were generally absent in healthy tissue and only detected within blood vessels. **(D)** GFP$^+$ neurons remained stable throughout 24 weeks of imaging, confirming long-term structural integrity.

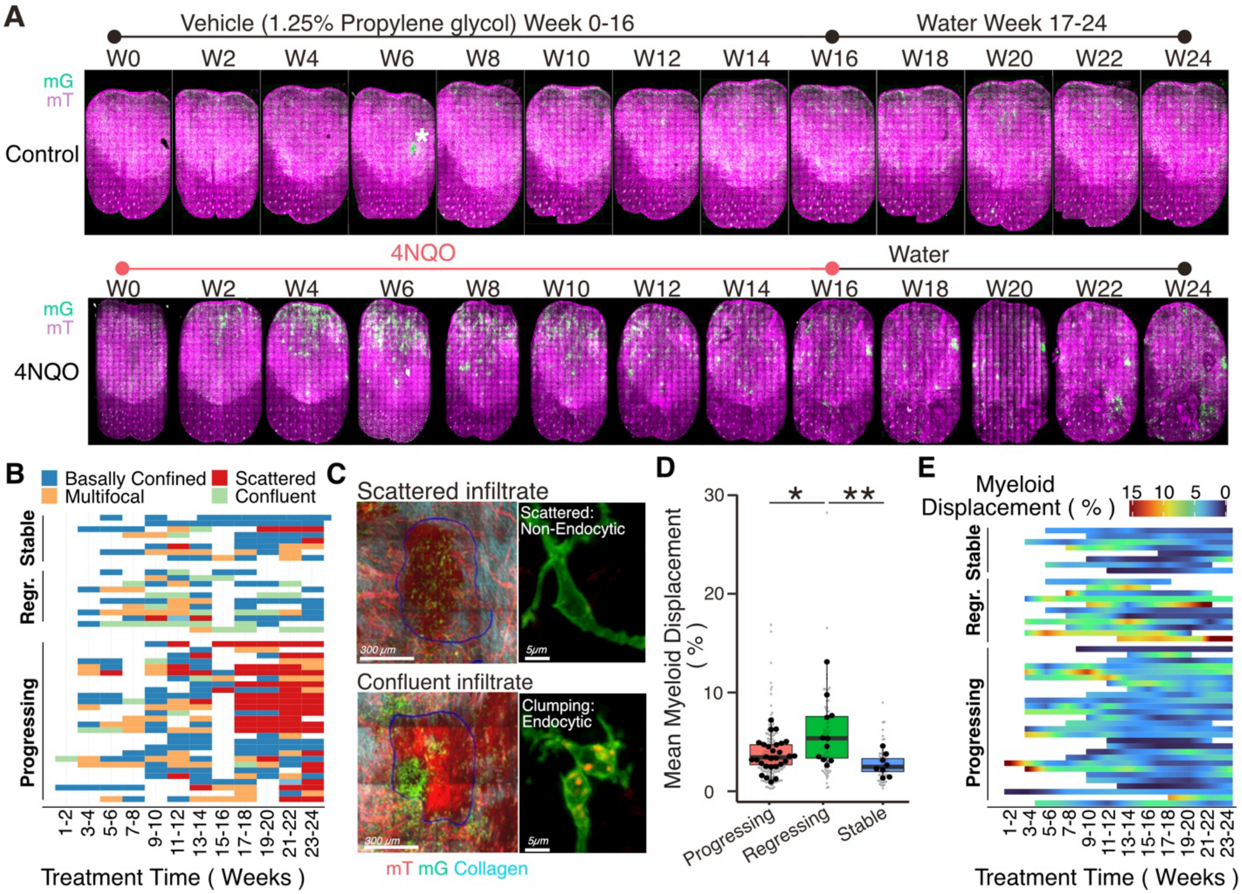


**Supplementary Figure 3 – Characterization of immune cells in the tongue 4NQO-treated mice (A)** Representative biweekly imaging results from a control mouse receiving vehicle only (1.25% propylene glycol in drinking water; top) and from a 4NQO-treated mouse (bottom). Two control mice were subjected to the same biweekly imaging schedule. Asterisk (*) indicates a transient wound in the control mouse at week 6 that was fully healed by week 8. **(B)** Bar graph illustrating the frequency of the four myeloid behavioral patterns described in Figure 2C in all individual lesions at all time points. Each row represents a single lesion. **(C)** Intravital images of clustered versus individual myeloid cells. Insets show single cells; clustered myeloid cells contain endocytosed TdTomato signal, indicative of active phagocytosis. **(D)** Boxplot of mean myeloid displacement (myeloid volume / lesion volume) across all time points for each lesion. Small dots represent individual lesion time points; large dots represent the mean per lesion. Differences between distributions were assessed by pairwise Wilcoxon tests. **(E)** As in (D), showing myeloid displacement stratified by treatment duration.

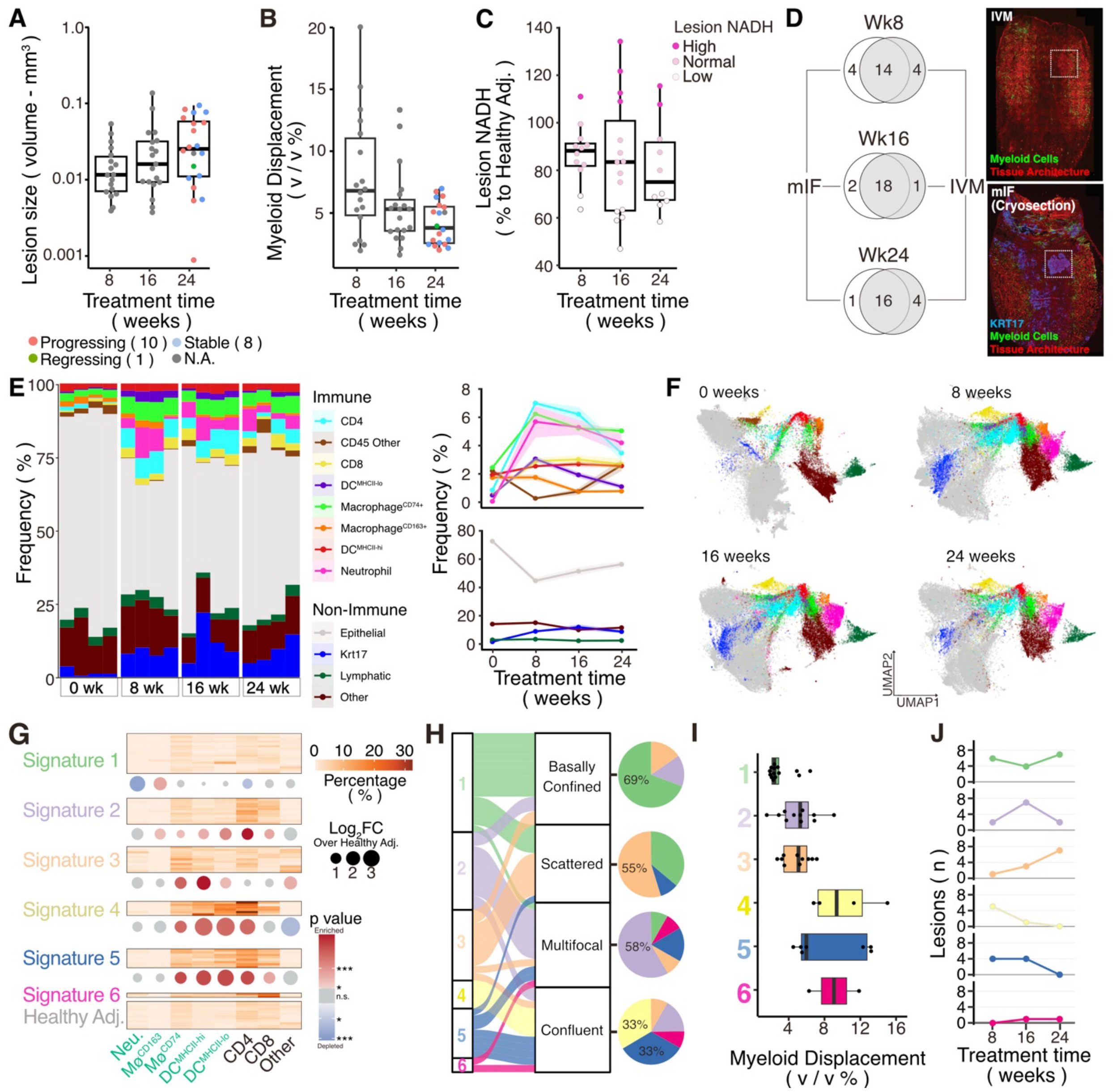

**Supplementary Figure 4 – Intravital summary statistics of lesions and multiplex immunofluorescence (mIF) within whole tongues of the correlative staining cohort.** **(A)** Boxplots showing average myeloid displacement for individual lesions in mice sacrificed at different treatment time points. **(B)** Boxplots summarizing myeloid displacement in lesions over the course of treatment. **(C)** Boxplots of average intratumoral NADH levels for individual lesions across time points, color-coded by NADH intensity. **(D)** Venn diagrams illustrating the overlap of lesions identified by mIF and intravital imaging at three different time points. **(E)** Stacked bar plots showing the frequency of each cell type per mouse (three sections analyzed per mouse, including both epithelium and lamina propria; four mice per time point). The accompanying line plots depict average frequencies of immune (top) and non-immune (bottom) cells, with ribbons representing mean ± SEM. **(F)** UMAP projections displaying the distribution of identified cell populations. **(G)** Heatmap showing the relative proportions of immune cell types within 62 lesions. Unsupervised clustering defined six immune signatures; dots represent fold enrichment (red) or depletion (blue) of immune subsets compared to healthy adjacent tissue, with color intensity corresponding to *p*-values from Mann–Whitney tests (gray = not significant). **(H)** Sankey plot linking immune signatures to intravital myeloid clustering behaviors. **(I)** Myeloid displacement values determined by IVM for lesions associated with each of the six immune signatures. **(J)** Line plot showing the frequency of lesions at different stages of 4NQO treatment.

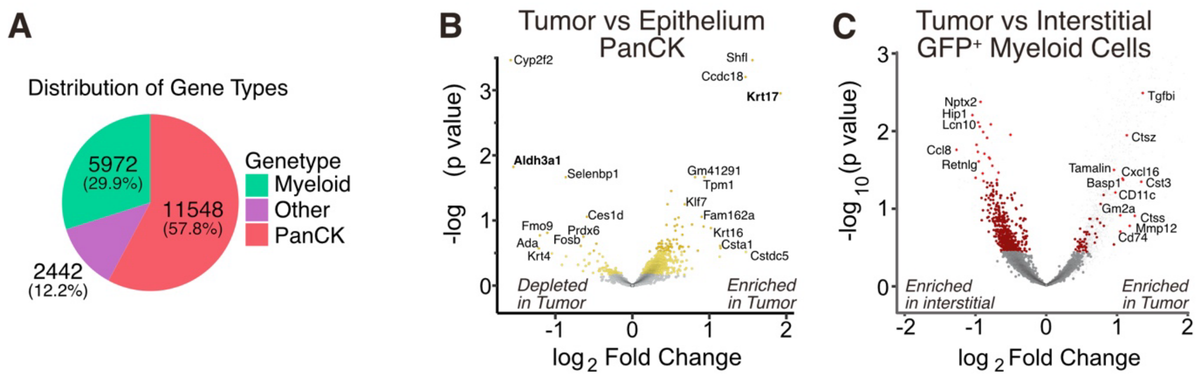


**Supplementary Figure 5 – GeoMx Digital Spatial Profiling reveals Antigen-presenting and myeloid-derived suppressor transcripts in endpoint lesions. (A)** Pie chart displaying the number of genes with the highest expression in either Myeloid ($GFP^+$), epithelial ($PanCK^+$) or Other ($GFP^-$ / $PanCK^-$) segments, defining gene transcripts associated with myeloid, epithelial or, other cells. **(B)** Volcano plot of transcriptomic counts from the GeoMx digital spatial Profiling from Epithelial ($PanCK^+$) segments within tumor regions compared to healthy adjacent epithelial tissue. Small transparent dots represent gene transcripts associated with non-epithelial cells. **(C)** Volcano plot as in B, comparing myeloid segments from tumor regions and isolated interstitial regions. A Linear Mixed Model was used to test difference in expression levels between lesions and healthy areas. Small transparent dots represent gene transcripts associated with non-Myeloid cells.

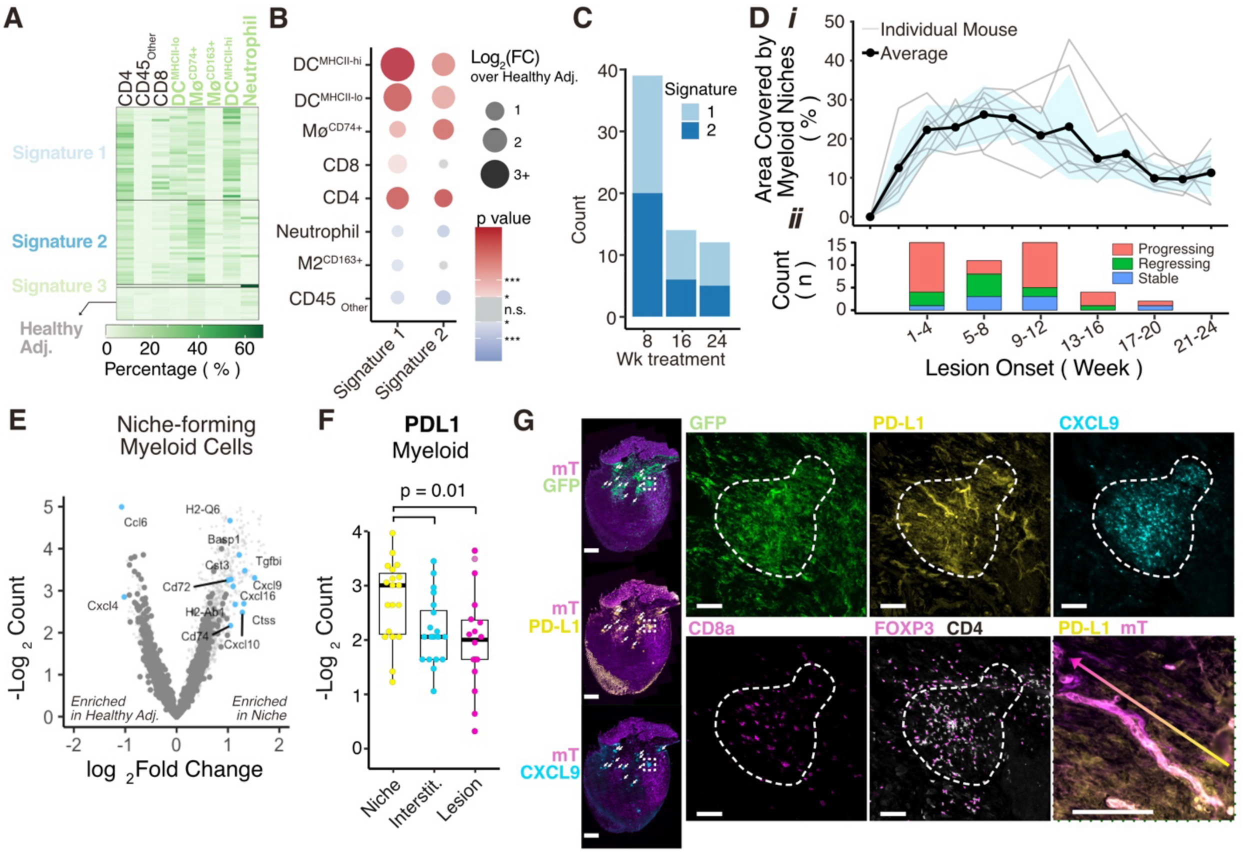


**Supplementary Figure 6 – Orphan clusters accumulate during early treatment timepoints, express PD-L1, and attract Tregs. (A)** Heatmap showing the relative percentage of each immune cell type within 66 clusters not directly associated with a lesion, analyzed by the multiplex IF workflow presented in Figure 3. Unsupervised clustering revealed 3 distinct groups, of which group 3 was only represented by a single cluster that consisted of neutrophils. **(B)** Dot plot showing the fold enrichment or depletion of immune subsets within the clusters over healthy adjacent background. **(C)** The distribution of the two cluster types across timepoints. **(D)** i) line plot showing the area of the papillae-less area of the ventral tongue that is covered by myeloid clusters. Data represents the mean of 7 mice +/- SD. ii) stacked bar graph showing the onset lesion divided by trajectory, and that the majority of lesions start in the early part of the plateau phase of myeloid clustering. **(E)** Volcano plot of transcriptomic counts from the GeoMx digital spatial Profiling from Myeloid cell (GFP+) segments within orphan cluster regions compared to healthy and non-clustering interstitial myeloid cells. **(F)** Boxplot showing transcriptomic results of PD-L1 in the myeloid Areas of Interest, segregated by region in which they were sampled, highlighting that niches have the highest expression of PD-L1. Statistical test: pairwise. **(G)** Immunofluorescence of sections of a tongue stained after 2weeks of 4NQO treatment, highlighting colocalization of CXCL9, PDL1, and myeloid niches. Scale bar = 1 mm. Inserts show a magnified view of a single cluster, highlighting CD8 T-cells and FOXP3+ Treg cells within the niche. Gradient arrow highlights that blood-vessels also express PD-L1 within myeloid clusters, which dissipates as the vessel continues out of the niche region. Scale bar = 100mm.

**Supplementary Video 1: Myeloid dynamics of scattered Infiltrates:**

Time-lapse imaging showing the dynamics of infiltrating myeloid cells with scattered behavior within a progressing lesion. Myeloid cells (mG: membrane GFP, green), collagen (SHG: second harmonic generation, cyan), and other cells (mT: membrane TdTomato). Total time: 10 min

**Supplementary Video 2: Myeloid dynamics of confluent infiltrates:**

Time-lapse imaging showing the dynamics of infiltrating myeloid cells with confluent behavior within a regressing lesion. Myeloid cells (mG: membrane GFP, green), collagen (SHG: second harmonic generation, cyan), and other cells (mT: membrane TdTomato). Total time: 10 min
